# Afadin mediates cadherin-catenin complex clustering on F-actin linked to cooperative binding and filament curvature

**DOI:** 10.1101/2024.10.08.617332

**Authors:** Rui Gong, Matthew J. Reynolds, Xiaoyu Sun, Gregory M. Alushin

## Abstract

The E-cadherin–β-catenin–αE-catenin (cadherin-catenin) complex couples the cytoskeletons of neighboring cells at adherens junctions (AJs) to mediate force transmission across epithelia. Mechanical force and auxiliary binding partners converge to stabilize the cadherin-catenin complex’s inherently weak binding to actin filaments (F-actin) through unclear mechanisms. Here we show that afadin’s coiled-coil (CC) domain and vinculin synergistically enhance the cadherin-catenin complex’s F-actin engagement. The cryo-EM structure of an E-cadherin–β-catenin–αE-catenin–vinculin–afadin-CC supra-complex bound to F-actin reveals that afadin-CC bridges adjacent αE-catenin actin-binding domains along the filament, stabilizing flexible αE-catenin segments implicated in mechanical regulation. These cooperative binding contacts promote the formation of supra-complex clusters along F-actin. Additionally, cryo-EM variability analysis links supra-complex binding along individual F-actin strands to nanoscale filament curvature, a deformation mode associated with cytoskeletal forces. Collectively, this work elucidates a mechanistic framework by which vinculin and afadin tune cadherin-catenin complex–cytoskeleton coupling to support AJ function across varying mechanical regimes.

## Introduction

Adherens junctions (AJs) are cell-cell adhesion complexes that transmit mechanical signals across epithelia to coordinate multicellular dynamics during development and tissue homeostasis^1–6^. AJ dysregulation leads to loss of cell-cell contact and epithelial-to-mesenchymal transition, hallmarks of cancer metastasis^7,8^. An interconnected protein network modulates the coupling strength between AJs and cytoskeletal actin filaments (F-actin) to support AJ function across the wide range of force regimes encountered in tissues^3,4,9^. The central component of this network is the heterotrimeric cadherin-catenin complex consisting of E-cadherin, β-catenin, and αE-catenin. E-cadherin is a transmembrane protein which directly mediates cell-cell contacts. The E-cadherin extracellular domains of two neighboring cells directly bind each other *in trans*, while their intracellular domains interact with β-catenin, which in turn associates with the actin-binding protein αE-catenin^9,10^. Multiple cadherin-catenin complexes further assemble into clusters within an AJ, mediated by *cis* interactions between E-cadherin extracellular domains within the same plasma membrane that are reinforced by cooperative binding of α-catenin to cytoplasmic F-actin^11–15^. This multi-layered network is critical for AJs to form mechanically robust connections between cells, yet it remains unclear how the interactions between its components are dynamically tuned to transmit and respond to cytoskeletal forces.

Force enhances F-actin binding by the cadherin-catenin complex, as well as its interactions with additional AJ proteins, to promote adhesion and facilitate mechanical signal transduction at AJs^6^. The cadherin-catenin complex in isolation is autoinhibited, displaying weak intrinsic binding affinity for F-actin and other adhesion partners^16–18^. Mechanical regulation is focused on α-catenin, whose C-terminal actin-binding domain (ABD) and central M domain undergo structural transitions associated with binding-partner engagement that can be modulated by force^19–21^. The complex forms a mechanically stabilized catch bond with F-actin in the presence of forces on the order of 10 pN. Mechanistically, this has been attributed to force-stabilized displacement of the first two alpha-helices (H0-H1) from the α-catenin ABD helical bundle, facilitating a torsional rearrangement of the bundle concomitant with strong F-actin binding^21,22^. Additionally, lower magnitude forces (∼1 pN) applied solely across F-actin enhance binding by the isolated αE-catenin ABD, which is mediated by the protein’s flexible 35 amino-acid C-terminal extension (CTE)^23^. Tension furthermore promotes unfurling of the αE-catenin M domain, facilitating binding by the αE-catenin structural homolog vinculin^19^. Vinculin engages the M1 subdomain through its N-terminal “head” domain, facilitating F-actin binding and catch bonding by its C-terminal “tail” ABD to reinforce cadherin-catenin complex–cytoskeleton coupling^16,18,24^. Additionally, the isolated vinculin head domain enhances the cadherin-catenin complex’s cooperative F-actin binding in the presence of mechanical load^25^. Despite substantial progress in establishing the biophysical mechanisms of this core network of force-modulated interactions, it remains largely unknown how the cadherin-catenin complex interfaces with additional critical adhesion proteins to maintain AJ mechanical integrity and function.

One such factor is afadin, a large (206 kDa) multidomain scaffolding protein with numerous AJ binding partners, including αE-catenin and F-actin^26–28^. Afadin knockout is embryonically lethal in the mouse, where defective cell-cell junctions fail to support development^29,30^. In human patients, dysregulated afadin expression has been associated with carcinogenesis and cancer metastasis in multiple tissue contexts, and somatic afadin loss-of-function mutations have recently been reported to drive a class of E-cadherin-positive metastatic breast cancer^31,32^. Afadin inactivation in cultured epithelial cells delays initial AJ formation without impacting the integrity of mature adhesions^33,34^, mirroring the mild phenotype of knocking out the *Drosophila* afadin homolog canoe in morphogenetically inactive tissues^35^. However, during periods of tissue rearrangement featuring elevated cellular contractility such as morphogenesis, afadin or canoe ablation causes defective AJ remodeling, impairing processes including apical constriction^35,36^; convergent extension^37,38^, wound healing^34^ and collective cell migration^39^. This supports a key role for afadin in reinforcing AJs under conditions of high tissue tension.

Although it is structurally divergent from vinculin, afadin features both αE-catenin and F-actin binding domains. This suggests that it might function in a conceptually analogous fashion to vinculin by reinforcing the interaction between the cadherin-catenin complex and F-actin^26,27^. However, canoe’s F-actin binding activity is dispensable for *Drosophila* morphogenesis^40^, challenging this model for afadin-mediated AJ stabilization. Recently, afadin’s short C-terminal coiled-coil region (residues 1393-1602, hereafter referred to as “afadin-CC”), which is required in canoe for morphogenesis^41^, has been reported to enhance F-actin binding by the cadherin-catenin complex through an unknown mechanism, providing an alternative explanation for afadin’s stabilizing effect^36^. Although both vinculin and afadin contribute to AJ formation and remodeling, it remains unclear how they coordinately tune the coupling strength between the cadherin-catenin complex and F-actin to adapt to various force conditions.

Here, we show that vinculin and afadin collaborate to promote a highly activated form of the cadherin-catenin complex, with afadin-CC physically engaging mechanically-regulated structural elements in αE-catenin to mediate cooperative F-actin binding. In biochemical assays, we find that vinculin and afadin-CC synergistically enhance the cadherin-catenin complex’s F-actin binding affinity. A 3.1 Å resolution cryogenic electron microscopy (cryo-EM) structure of the five-component mammalian E-cadherin–β-catenin–αE-catenin–vinculin–afadin-CC supra-complex bound to F-actin resolves extensive contacts between afadin-CC and the αE-catenin ABD. Afadin-CC directly promotes cooperativity by bridging two adjacent ABDs, binding and stabilizing both the displaced helix H1 and the CTE. This induces folding of the entire CTE on the F-actin surface, which mediates additional F-actin and intra-ABD contacts, resulting in an extensive interaction network that tightly anchors neighboring cadherin-catenin complexes. Consistently, total internal reflection fluorescence (TIRF) studies show that the afadin-CC substantially enhances the cadherin-catenin complex’s cooperative F-actin binding, promoting the formation of supra-complex clusters along filaments. Furthermore, ablation of specific afadin-CC–αE-catenin ABD contacts eliminates afadin-CC’s stimulation of cadherin-catenin complex F-actin binding *in vitro* and disrupts AJ organization in cultured cells. From our cryo-EM dataset, we also obtained a structure featuring the supra-complex asymmetrically bound along one F-actin strand, which variability analysis links to F-actin curvature. This suggests afadin’s enhancement of cooperative F-actin binding by the cadherin-catenin complex is likely additionally promoted by nanoscale mechanical deformation of F-actin by cytoskeletal forces. Collectively, these data supporting a model in which afadin enhances the cadherin-catenin complex’s cytoskeletal engagement by stabilizing a network of binding interactions on the F-actin surface, simultaneously bolstering the effects of mechanical force and binding partners such as vinculin.

## Results

### Reciprocally stimulated F-actin binding by the cadherin-catenin complex and vinculin

To probe how the interplay of multiple actin-binding adhesion proteins can modulate AJ– cytoskeleton coupling strength, we first examined the cadherin-catenin complex and vinculin in F-actin co-sedimentation assays. In isolation, both factors have been reported to adopt a closed conformation, displaying minimal binding to each other and to F-actin^17,20,42^. Consistently, we find that wild-type (WT) full-length vinculin and a minimal soluble ternary cadherin-cadherin complex (hereafter “Eβα”) which includes the intracellular β-catenin binding region of E-cadherin, full length β-catenin and full-length αE-catenin, each individually exhibited weak F-actin binding (Figures 1A, S1A and S1D). To test whether binding between vinculin and the cadherin-catenin complex mutually influences their F-actin engagement, we introduced mutations previously reported to reduce autoinhibitory interactions in vinculin^43^ (D974A/K975A/R976A/R978A, “vinculin(T12)”) and αE-catenin^44^ ( M319G/R326E/R551E, “α(CA)”) in the context of the minimal Eβα trimer. Unexpectedly, under our assay conditions both mutants still displayed weak F-actin binding in isolation, comparable to that of the respective wild-type proteins, implying that their ABDs are still substantially autoinhibited (Figures 1B and 1D). However, we find that the mutant proteins strongly interact, allowing us to reconstitute a stoichiometric Eβα(CA)-vinculin(T12) tetramer that exhibited significantly increased F-actin binding (Figures 1B, 1D and S1E).

**Figure 1.**
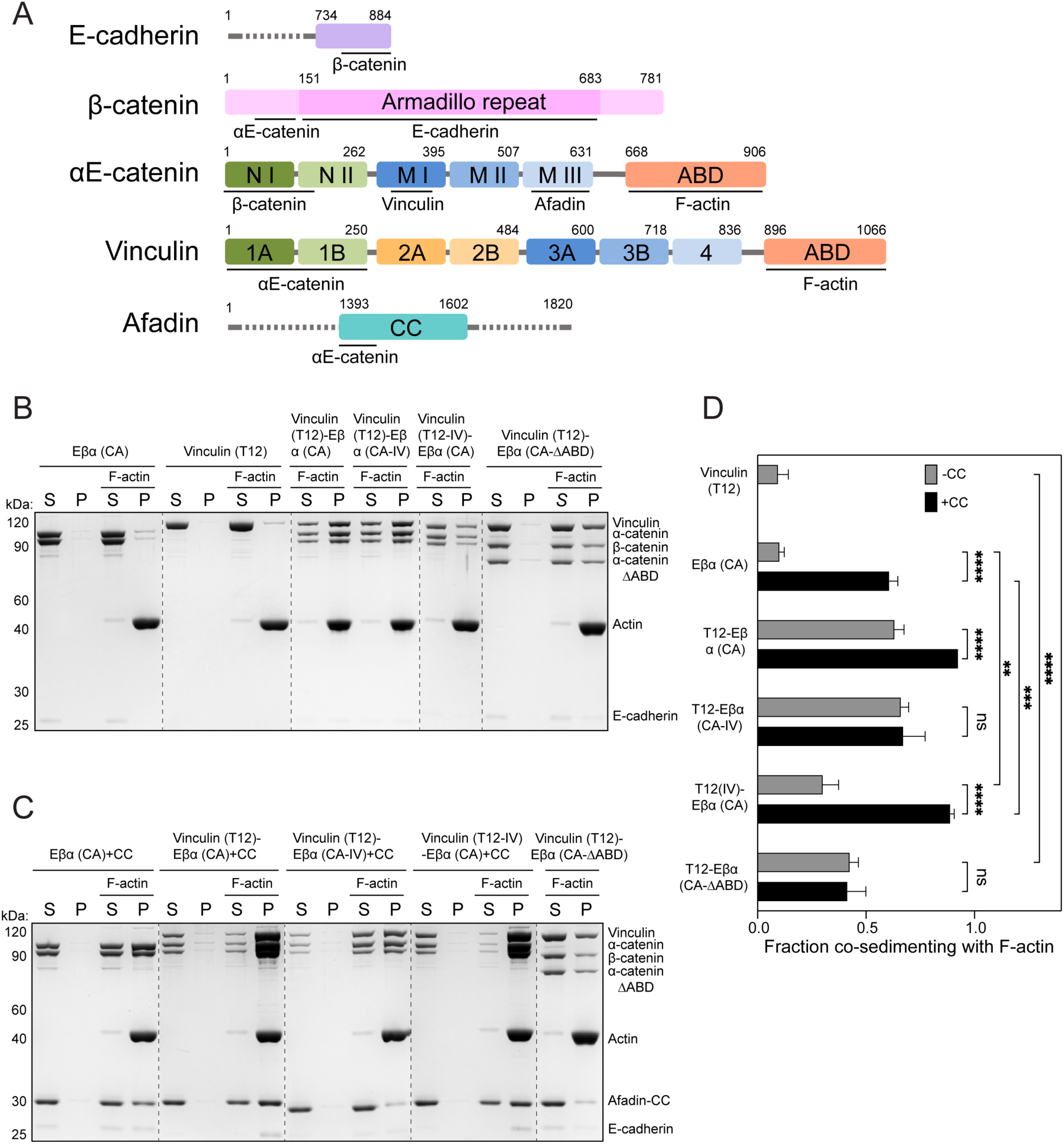
Vinculin and afadin synergistically enhance the cadherin-catenin complex’s F-actin binding. (A) Domain organization of E-cadherin, β-catenin, αE-catenin, vinculin, and afadin. Domains used in this study are colored and sequence boundaries are indicated. ABD, actin binding domain; CC, coiled coil. (B-C) Co-sedimentation assays showing the binding of indicated proteins and protein complexes to F-actin. S, supernatant; P, pellet. (D) Quantification of the fraction of indicated proteins or protein complexes which co-sedimented with F-actin in the pellet. Measurements were corrected by subtracting the amount of protein which sedimented in the absence of F-actin. Data are presented as mean ± SD of three independent experiments. Conditions were compared using two-way ANOVA with Tukey’s multiple comparison test: NS, p ≥ 0.05; *p < 0.05; **p < 0.01; ***p < 0.001; ****p < 0.0001. Dotted lines indicate stitching interfaces between gels.

We next assessed the contributions of the vinculin and αE-catenin ABDs to this enhanced F-actin binding activity. We first validated point mutations previously reported to reduce F-actin binding in vinculin (I997A/V1001A, “vinculin (IV)”) and αE-catenin (I792A/V796A, “α(IV)”) by examining their isolated ABDs^45,46^. Consistent with previous studies, the IV mutations reduced F-actin binding by the vinculin ABD to undetectable levels in our assay, and substantially attenuated F-actin binding by the αE-catenin ABD (Figures S1C and S1D). We then examined these mutants in the context of the higher-order complex. The IV mutations in vinculin (Eβα(CA)–vinculin(T12-IV)) markedly reduced F-actin binding of the tetramer, which nevertheless bound better than the minimal cadherin-catenin complex trimer (Eβα(CA); Figures 1B and 1C). Conversely, these mutations in αE-catenin (Eβα(CA-IV)–vinculin(T12)) had only a marginal impact on the tetramer’s F-actin engagement (Figures 1B and 1C). As the IV mutations reduced but did not eliminate actin-binding by the αE-catenin ABD, we also examined a complex where this domain was completely deleted (Eβα(CA-ΔABD)–vinculin(T12)). Removing the αE-catenin ABD substantially attenuated F-actin binding, yet this complex nevertheless bound more tightly than vinculin(T12) (Figures 1B and 1C). Collectively, these data show that vinculin and the cadherin-catenin complex reciprocally enhance each other’s F-actin engagement. As the vinculin–cadherin-catenin complex interaction is known to occur through the vinculin Vh domain–αE-catenin M1 domain interface (Figure 1A)^20^, enhanced F-actin binding is likely mediated by an allosteric effect that releases both ABDs. While we find that the vinculin ABD is the primary actin-binding element of this complex, consistent with a recent report^47^, our data suggest that the αE-catenin ABD nevertheless contributes to its overall F-actin engagement.

### Afadin and vinculin synergistically enhance F-actin binding by the cadherin-catenin complex

We next examined the effects of afadin on our reconstituted AJ complexes. The afadin-CC domain has previously been shown to enhance the F-actin binding of a chimerically fused β-catenin-αE-catenin dimer^36^. Consistently, we find that the afadin-CC also markedly stimulates F-actin binding by the minimal constitutively active trimeric complex (Eβα(CA); Figures 1C and 1D). However, this effect was absent for the wild-type trimer (Eβα), and a trimer featuring IV mutations in the αE-catenin ABD (Eβα(CA-IV); Figures S1B and S1D), suggesting afadin-CC stimulates αE-catenin ABD release to potentiate its F-actin binding. Consistent with this model, pulldown assays with αE-catenin fragments confirm a strong interaction specifically between afadin-CC and the M3 domain of αE-catenin (Figure S2A)^48^. The distinct binding sites of vinculin (M1) and afadin (M3) on αE-catenin prompted us to investigate whether they can coincidentally engage the cadherin-catenin complex. In pulldown assays, we indeed find that afadin-CC and activated vinculin simultaneously bind the minimal constitutively active trimer to form a stable pentameric complex (Eβα(CA)–vinculin(T12)–afadin-CC; Figures S2B and S2C). This pentamer binds F-actin even more robustly than the tetrameric vinculin–cadherin-catenin complex (Eβα(CA)–vinculin(T12); Figures 1C and 1D), indicating that vinculin and afadin can synergize to enhance the cadherin-catenin complex’s F-actin engagement.

In striking contrast to the tetrameric cadherin-catenin-vinculin complex (Eβα(CA)–vinculin(T12-IV)), disrupting vinculin’s F-actin binding activity had minimal impact on the stimulating effect of afadin-CC in the pentameric complex (Eβα(CA)–vinculin(T12-IV)–afadin-CC; Figures 1C and 1D), suggesting this assembly instead primarily engages F-actin through the αE-catenin ABD. Moreover, vinculin featuring IV mutations (Eβα(CA)–vinculin(T12-IV)–afadin-CC) still stimulated the cadherin-catenin complex’s F-actin binding to a greater degree than afadin-CC alone (Eβα(CA)–afadin-CC; Figures 1C and 1D), indicating that afadin and vinculin converge to enhance F-actin binding through αE-catenin in the context of the pentameric complex. Consistently, complexes featuring lesions to the αE-catenin ABD (either IV mutations, Eβα(CA-IV)–vinculin(T12)–afadin-CC, or ABD deletion, Eβα(CA-ΔABD)–vinculin(T12)–afadin-CC) displayed substantial reductions in F-actin binding (Figures 1C and 1D). However, these complexes nevertheless bound F-actin to a similar degree as the corresponding tetrameric complexes in the absence of afadin-CC (Eβα(CA-IV)–vinculin(T12) / Eβα(CA-ΔABD)– vinculin(T12); Figures 1C and 1D), indicating that the vinculin ABD remains active in the pentamer, albeit making a diminished contribution to overall F-actin engagement. Collectively, these data indicate that afadin-CC and vinculin can simultaneously bind the cadherin-catenin complex to synergistically enhance its cytoskeletal association, primarily by stimulating F-actin binding through the αE-catenin ABD.

### Structure of a cadherin-catenin–vinculin–afadin-CC supra-complex bound to F-actin

To investigate the mechanisms by which AJ partners enhance the cadherin-catenin complex’s F-actin binding, we performed cryo-EM structural studies. To minimize heterogeneity at the F-actin interface, we focused on the pentameric complex featuring IV mutations in vinculin (Eβα(CA)-vinculin(T12-IV)-afadin-CC) bound to F-actin. Previous studies have suggested a highly flexible linker between the αE-catenin ABD and the M domain^49,50^. Consistently, in cryo-electron tomograms, we observed extended fibrous densities connected to actin filaments (Video S1), consistent with the 344 kDa mass of the complex, which appear to have no fixed orientation relative to the filament axis. We nevertheless pursued single particle cryo-EM, reasoning we might observe modulation of the αE-catenin ABD–F-actin interface that contributes to enhanced binding in the presence of afadin-CC.

Despite a substantial excess of pentamer (2.5 μM) versus F-actin (0.6 μM), we observed only partial decoration (Figure S3), potentially due to crowding effects imposed by the large flexible mass of the complex. This, along with the strong signal from these flexible regions in the raw micrographs, presented a challenge for traditional template-based particle-picking. We therefore employed a neural network-based approach (Methods). Following 2D classification, we observed only a small additional ordered density adjacent to F-actin (Figure S3), consistent with our tomography studies suggesting the vast majority of the complex is flexibly tethered to the filament (Video S1). Two distinct categories of 2D classes included this extra density. One features straight F-actin with decoration along both strands, consistent with prior structural studies of the isolated αE-catenin ABD bound to F-actin^23,51^, while the other unexpectedly features curved F-actin with decoration along only one strand (Figure S3).

To obtain a detailed view of the F-actin binding interface, we first focused on the well-ordered straight filament classes, which we reasoned would yield the highest-resolution 3D reconstruction. Analysis of these segments produced a final density map at 3.1 Å resolution (Methods; Figure 2A, S4A and S4B). While this map predominantly resembled previously reported reconstructions of the isolated αE-catenin ABD bound F-actin^23,51^, it featured three additional densities. The first is a coiled-coil density bridging two adjacent αE-catenin ABDs, which we identified as residues 1508-1578 of the afadin-CC using an AlphaFold2 predicted structure of this segment (Figures S4C-S4E)^52^. The second density extended from the previously resolved proximal portion of αE-catenin’s CTE (residues 844-871, which we here refer to as “CTE-N”), indicating folding of the disordered distal portion of the CTE (“CTE-C”, residues 872-906; Figures S4C-S4E). High-quality density in two regions (residues 881-884 and 891-903) enabled us to model the entire CTE except for the final 3 residues (Figure S4E). The third and final extra density, a helical segment tucked alongside the afadin coiled-coil, was unambiguously assigned as residues 690-706 (the carboxyl-terminal half of H1) of the αE-catenin ABD, owing to its connection to H2 (Figures S4C-S4E). Our final refined atomic model contains the entire sequence of actin except for the first 4 residues, residues 690-903 of αE-catenin, and residues 1508-1578 of afadin.

**Figure 2.**
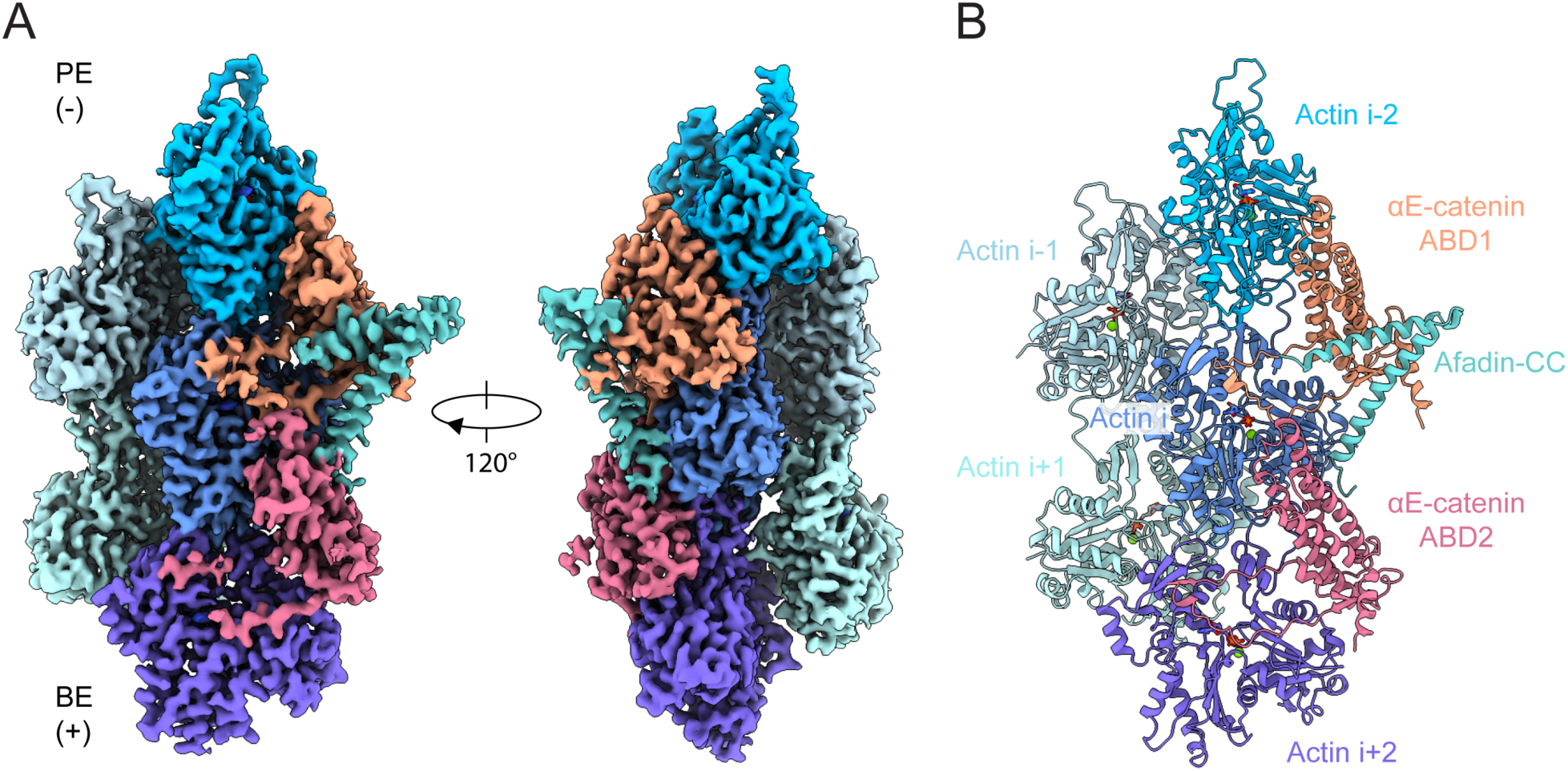
Cryo-EM structure of pentamer bound F-actin, visualizing αE-catenin ABD stabilization by afadin-CC. (A) 3.1 Å resolution cryo-EM density map, which resolves two αE-catenin ABDs and one afadin-CC bound to F-actin. BE, barbed end; PE, pointed end. (B) Ribbon representation of the associated atomic model.

In the structure, each αE-catenin ABD interacts with two neighboring actin subunits along the same strand, while each actin subunit contacts two adjacent ABDs (Figure 2B). The two arms of the “V” shaped afadin coiled-coil point towards the central axis of the filament, with each arm engaging one of the two neighboring ABDs. At each interface, the C-terminal arm of the coiled-coil makes extensive contacts with the ABD towards the minus (“pointed”) end of the filament (ABD1), while the N-terminal arm makes more limited contacts with the ABD towards the plus (“barbed”) end (ABD2; Figure 2B).

### Afadin stabilizes flexible elements of the αE-catenin ABD implicated in force sensitivity

Previous structural studies of the αE-catenin ABD have shown that substantial conformational remodeling accompanies its F-actin engagement^23,51^. This includes an order-to-disorder transition of H0-H1, a torsional repacking of the H2-H5 helical bundle, and a shift of the proximal CTE-N, which collectively facilitate high-affinity binding to the filament (Figures 3A, 3B, 3D and Video S2). Functional studies have suggested that H0-H1 release is promoted by tension to mediate catch-bond formation^22,45^, while CTE-C is required for force-activated F-actin binding when load is applied across the filament^23^. Superimposing the αE-catenin ABD from our supra-complex structure on our previous structure of the isolated ABD bound to F-actin shows nearly identical conformations of H2-H5 (residues 707-843, RMSD of 0.466 Å for 118 aligned Cα atoms) and the F-actin binding region of CTE-N (residues 863-871, RMSD of 0.290 Å for 9 aligned Cα atoms; Figure 3E). Modulation of this interface is therefore unlikely to contribute to the supra-complex’s enhanced F-actin affinity. However, the afadin coil-coil induces substantial remodeling of H1, reconfiguration of the proximal region of CTE-N, and the complete refolding of CTE-C to mediate specific F-actin and intra-ABD contacts, suggesting afadin’s engagement of these flexible elements potently stabilizes F-actin binding (Figures 3C and 3E).

**Figure 3.**
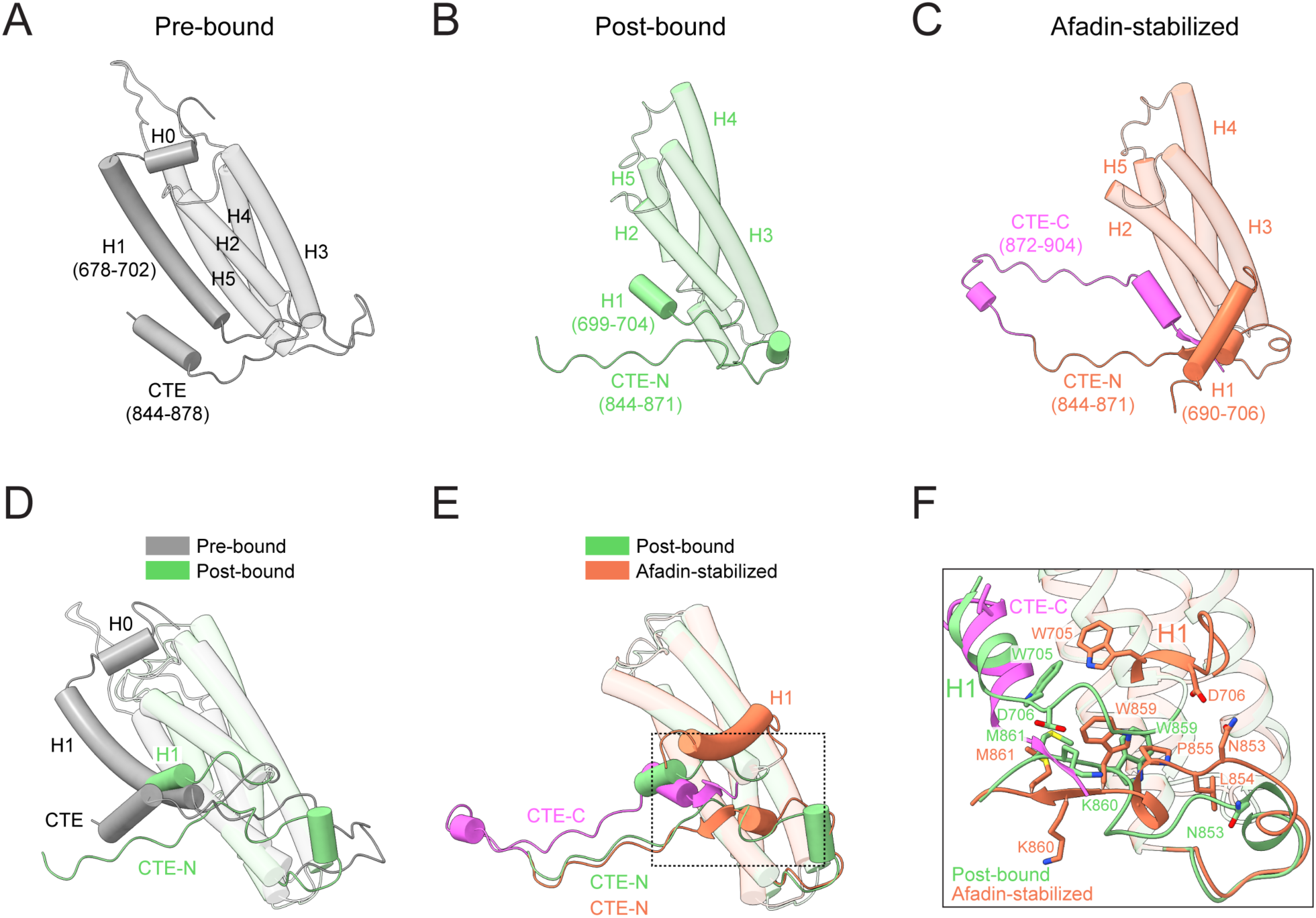
Structural remodeling of the αE-catenin ABD evoked by afadin-CC. (A-C) Structures of the αE-catenin ABD in three different conformational states. Pre-bound, crystal structure of αE-catenin (PDB: 4IGG chain B); post-bound, cryo-EM structure of αE-catenin ABD alone bound to F-actin (PDB: 6UPV); afadin-stabilized, this study. CTE, C-terminal extension. The refolded CTE-C is highlighted in magenta. (D) Superposition of the αE-catenin ABD in the pre-bound and post-bound states. (E) Superposition of the αE-catenin ABD in the post-bound and afadin-stabilized states. (F) Detail view (boxed in E) highlighting αE-catenin ABD H1 and CTE structural rearrangements evoked by afadin-CC engagement.

In previous structures of the isolated αE-catenin ABD in the absence of F-actin (the “pre-bound” state), the fully folded H1 packs against H2 and H5 (Figure 3A and Video S2)^50^. In structures of the isolated αE-catenin ABD bound to F-actin, a short stub of H1 remains docked on the helical bundle upon F-actin binding (the “post-bound” state; Figure 3B and Video S2). In striking contrast, H1 is fully displaced in the afadin stabilized state (Figure 3C and Video S3). H1 from ABD1 is repositioned to bind beneath the central joint of the afadin coiled-coil, where residues 694-698 become reordered, forming interactions that mutually stabilize afadin and αE-catenin. Furthermore, residues W705 and D706 immediately C-terminal to H1 rearrange, incorporating to extend the helix (Figures 3C and 3F).

While CTE-C was unresolved in both the pre-bound and post-bound states of the isolated αE-catenin ABD, afadin strikingly stabilizes refolding of the entire αE-catenin CTE on the F-actin surface, pinioning it beneath the C-terminal arm of the coiled-coil (Figures 3C, S4D and Video S3). The ordered CTE-C of ABD1 first extends upward through a flexible linker (residues 872-880), followed by a patch of well-resolved residues (881-884) which interface with subdomain 4 of actin i. It then turns toward the H2-H5 helical bundle of ABD1 through another linker (residues 885-892). The carboxyl-terminal residues 893-898 refold into a short α-helix, which sits in a groove vacated by complete displacement of H1 (Figures 3E and 3F). This is followed by a short β-strand (residues 900-902), which inserts into a pocket generated by remodeling of residues 859-861, where it is surrounded by the C-terminal coil of afadin-CC, as well as H1, H2, H5 and CTE-N of ABD1 (Figures 3E and 3F). To accommodate the newly inserted CTE-C, residues 853-861 of the proximal CTE-N undergo substantial remodeling. Residues 859-861 shift downward from the helical bundle and fold into a short β-strand, forming a stabilizing β-sheet with residues 900-902. Reciprocally, residues 853-855 move upward to fill the space left by the displacement of the bulky residue W859, which was found to undergo minimal rearrangements in the isolated αE-catenin ABD–F-actin structure, where it restricts rearrangements of CTE-N (Figures 3E and 3F)^23,51^.

### Afadin–αE-catenin interfaces mediating supra-complex binding to F-actin

Afadin binding and the concomitant structural rearrangements of the αE-catenin ABD establish multiple interfaces likely to strengthen the interaction between the cadherin-catenin complex and F-actin. At the N terminus of the first afadin coil, a short segment comprising residues 1509-1518 forms a tripartite interface with ABD2 and actin subunit i (Figure 4A and 4B). This interface features a combination of hydrogen bonds, Van der Waals contacts, and salt bridges. Notably, W1514 from afadin embeds into a groove formed by H3 and H4 of ABD2, while R1516 hydrogen bonds with A22 and D24 of actin i. In addition, D1517 forms a salt bridge with αE-catenin K795.

**Figure 4.**
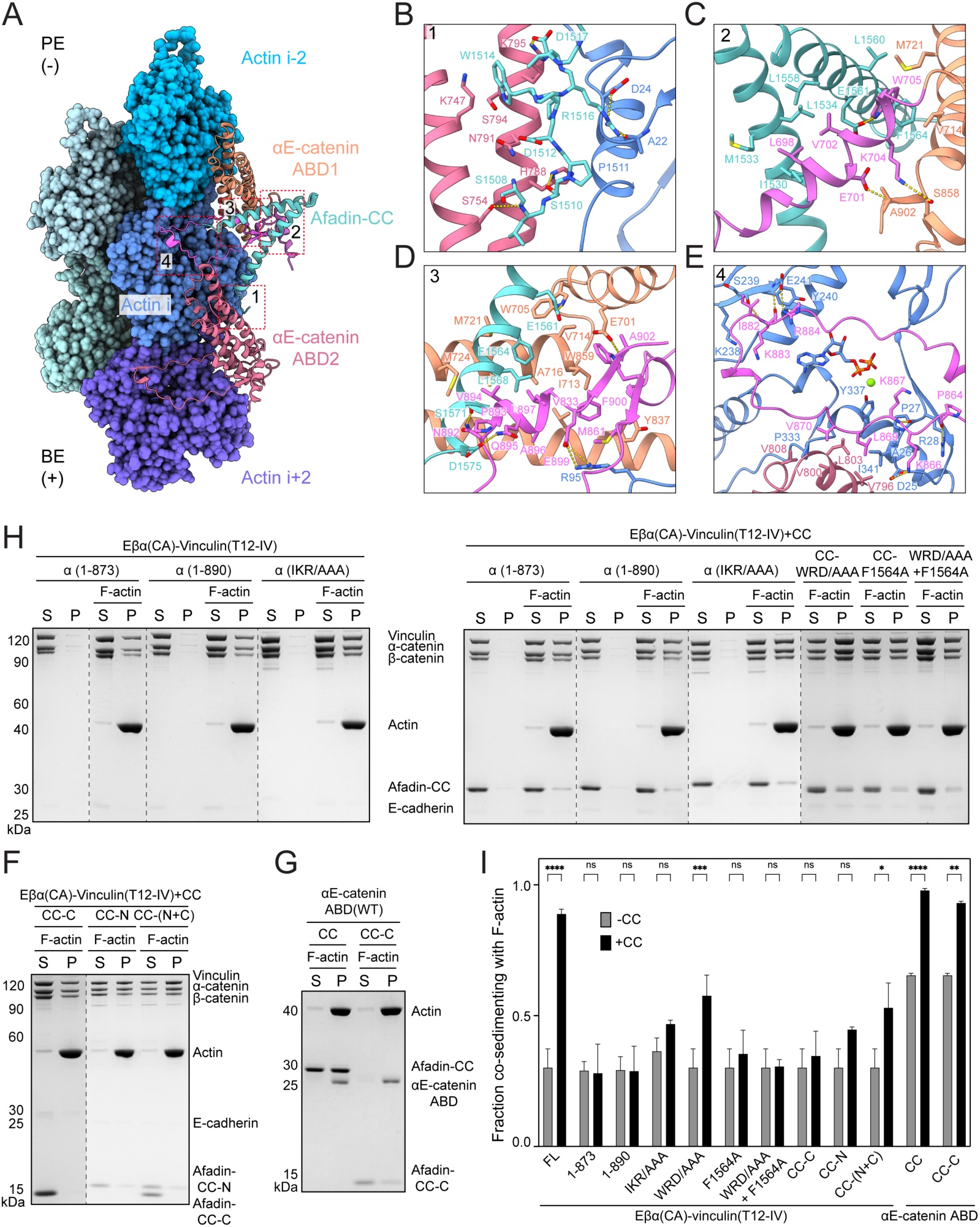
Afadin-CC engagement mediates enhanced binding of the αE-catenin ABD to F-actin. (A) Overview of the tripartite interface. BE, barbed end; PE, pointed end. (B) Contacts between actin subunit i, αE-catenin ABD2, and afadin-CC’s N-terminal coil. (C) Contacts at the interface between the displaced H1 and H2 of αE-catenin ABD1 and afadin-CC. (D) Interaction network formed by afadin-CC’s C-terminal coil, αE-catenin ABD1, and actin subunit i. (E) Contacts between the CTE of αE-catenin ABD1, actin subunit i, and H4 of αE-catenin ABD2. (F-G) Co-sedimentation assays analyzing the effects of afadin-CC fragments on F-actin binding by the tetrameric complex (F) and the isolated αE-catenin ABD (G). (H) Co-sedimentation assays analyzing the contributions of specific interfaces to afadin-CC’s stimulation of F-actin binding by the tetramer. (I) Quantification of F-H. Data are presented as mean ± SD of three independent experiments. Conditions were compared by two-way ANOVA with Tukey’s multiple comparison test: NS, p ≥ 0.05; *p < 0.05; **p < 0.01; ***p < 0.001; ****p < 0.0001. Dotted lines indicate stitching interfaces between gels.

The rearranged H1 helix of ABD1 is sandwiched by the coiled-coil of afadin and the CTE-N of ABD1. The interaction between H1 and the coiled-coil is mediated by a predominantly hydrophobic interface. Afadin-induced repositioning of H1 places W705 in a newly formed hydrophobic groove between the second coil of afadin and H5 of ABD1. In addition, several hydrophilic H1 residues on the opposite side of the helix make electrostatic interactions with both CTE-N and CTE-C (Figure 4C).

The folded short α-helix of CTE-C (893-898) is sandwiched by the C-terminal coil of afadin-CC and H2 / H5 of ABD1 (Figure 4D). The following short β-strand (900-902) and the resulting β-sheet further packs onto the sandwich module, creating an intertwined interaction network primarily driven by hydrophobic interactions. Hydrophobic residues in the CTE-C, including V894, A896, L897, and F900, deeply penetrate into the hydrophobic groove created by H2 and H5, which is further fortified by F1564 and L1568 from afadin, as well as W859 and M861 from CTE-N. In addition to these hydrophobic interactions, the rearranged CTE-C also forms several electrostatic interactions with afadin and F-actin (Figures 4C and 4D).

The conformation of CTE-N residues 862-871 remains essentially unaltered after afadin engagement versus the post-bound state (Figure 3E). The proximal segment electrostatically interacts with actin i, while the distal segment sits in a hydrophobic cleft cradled by actin i and ABD2, where previously reported direct contacts with ABD2 are clearly resolved in the supra-complex structure (Figure 4E)^23,51^. Beyond this region, a flexible linker enables a patch of previously disordered residues (882-884) to align in parallel with a β-strand from subdomain 4 of actin i, forming to our knowledge a previously unreported interface between αE-catenin and F-actin (Figures 4A and 4E). The contacts feature a salt bridge between αE-catenin R884 and actin E241, backbone hydrogen bonding between αE-catenin K883 and actin S239 and E241, as well as Van der Waals contacts involving αE-catenin I882 (Figure 4E).

Collectively, this extensive network of interactions likely directly stabilizes the cadherin-catenin complex’s F-actin binding in the presence of afadin.

### The αE-catenin ABD-afadin-CC interface is required for proper actomyosin organizations at AJs

Although we visualized extensive contacts between residues 1510-1602 of the afadin-CC domain (hereafter referred to as afadin-CC-C) and the αE-catenin ABD, residues 1393-1510 (afadin-CC-N) were not resolved in our structure. As this sequence is responsible for binding to αE-catenin’s M3 domain (Figure S2D)^48^, we hypothesized that it could also contribute to activation by promoting ABD release, in a manner conceptually analogous to the vinculin Vh domain. We thus sought to determine the interplay of these two afadin-CC segments in enhancing the cadherin-catenin complex’s F-actin engagement using co-sedimentation assays. Neither isolated afadin-CC-N nor afadin-CC-C alone had a stimulatory effect on F-actin binding by the tetramer (Eβα(CA)-vinculin (T12-IV); Figure 4F and 4J). This suggests that afadin-CC-C’s direct stabilization of the F-actin binding interface can only occur if αE-catenin ABD release is promoted by afadin-CC-N. In agreement with this model, both afadin-CC and afadin-CC-C potently stimulated F-actin binding by the isolated αE-catenin ABD, which completely lacks autoinhibition (Figure 4G and 4J). Furthermore, a mixture of afadin-CC-N and afadin-CC-C substantially increased F-actin binding by the tetramer, consistent with our proposed hierarchy of activities, albeit less potently than the intact afadin-CC (Figure 4F and 4J). This suggests the physical linkage of these two segments enhances afadin-CC’s overall effect on stimulated F-actin binding, potentially by increasing the local concentration of afadin-CC-C in the vicinity of αE-catenin ABD activated by afadin-CC-N.

We next dissected the afadin-CC-C / αE-catenin ABD interfaces we visualized by designing mutations to selectively disrupt them in the context of the pentamer featuring IV mutations in vinculin (Eβα(CA)-vinculin (T12-IV)-afadin-CC) and performing co-sedimentation assays. In the refolded αE-catenin CTE-C region, truncation of either the entire CTE-C (874-906) or the C-terminal short α-helix and β-strand (891-906) did not impact the F-actin binding of the tetrameric complex (Eβα(CA)– vinculin(T12-IV)) in the absence of afadin-CC, but these lesions completely abolished afadin-CC’s stimulatory effect (Figure 4H, 4I and 4J). Moreover, mutations disrupting the interface between the proximal CTE-C (I882A/K883A/R884A, “IKR/AAA”) and F-actin led to a significant reduction in stimulation (Figure 4H, 4I, and 4J). These results together demonstrate that both the actin and afadin binding regions of αE-catenin’s CTE-C are indispensable for enhancement of F-actin binding by afadin. While CTE-C is highly conserved across metazoans and among different α-catenin isoforms, an equivalent segment is absent in vinculin (Figure S5A). Thus, the CTE-C segment likely plays a conserved role in specifically mediating enhancement of cadherin-catenin complex–F-actin binding through the α-catenin ABD.

In the first coil of afadin-CC, alanine substitutions of the key ABD2 interacting residues W1514, R1516 and D1517 (W1514A/R1516A/D1517A, “WRD/AAA”) substantially decreased F-actin binding stimulation (Figure 4I and 4J). Similarly, a single alanine substitution of F1564 within the second afadin-CC coil, which is involved in the core of hydrophobic interactions with ABD1, nearly abolished afadin-CC’s stimulatory effect (Figure 4I and 4J). Combining these mutations reduced afadin-CC’s stimulatory effect to an undetectable level (Figure 4I and 4J). These findings suggest that afadin’s interaction with both αE-catenin ABDs is important for its functional modulation of the F-actin binding interface, with ABD1 contacts playing a more pronounced role. Notably, the segments of the afadin coiled-coil observed in our structure are highly conserved from *Drosophila* to humans (Figure S5B), suggesting it plays a similar role in stabilizing the αE-catenin–F-actin interaction across species.

In EpH4 mouse mammary epithelial cells, afadin-CC has been shown to be necessary and sufficient for proper actin-myosin (actomyosin) bundle organization at AJs^36^. In wild-type cells, afadin co-localizes with E-cadherin at AJs. The actomyosin network, as indicated by F-actin staining, forms a distinct, thick line along the junctions between adjacent cells, implying tight coupling between the cadherin-catenin complex and junctional actomyosin (Figure 5). However, in afadin knockout cells, while E-cadherin maintains its localization to the plasma membrane, the F-actin signal broadens, indicating weakened cytoskeleton-AJ coupling (Figures 5 and S6). Consistent with previous studies^36^, re-expressing afadin-CC alone is sufficient to restore the single thick band of F-actin and its co-localization with E-cadherin (Figures 5 and S6). However, expression of mutant afadin-CC (W1514A/R1516A/D1517A/F1564K) failed to restore proper actomyosin architecture at AJs, consistent with this construct’s inability to enhance the cadherin-catenin complex’s binding to F-actin *in vitro* (Figures 4I, 5, and S6). This suggests direct interactions between the afadin coiled-coil and αE-catenin’s ABD are necessary for proper cellular actomyosin organizations at AJs.

**Figure 5.**
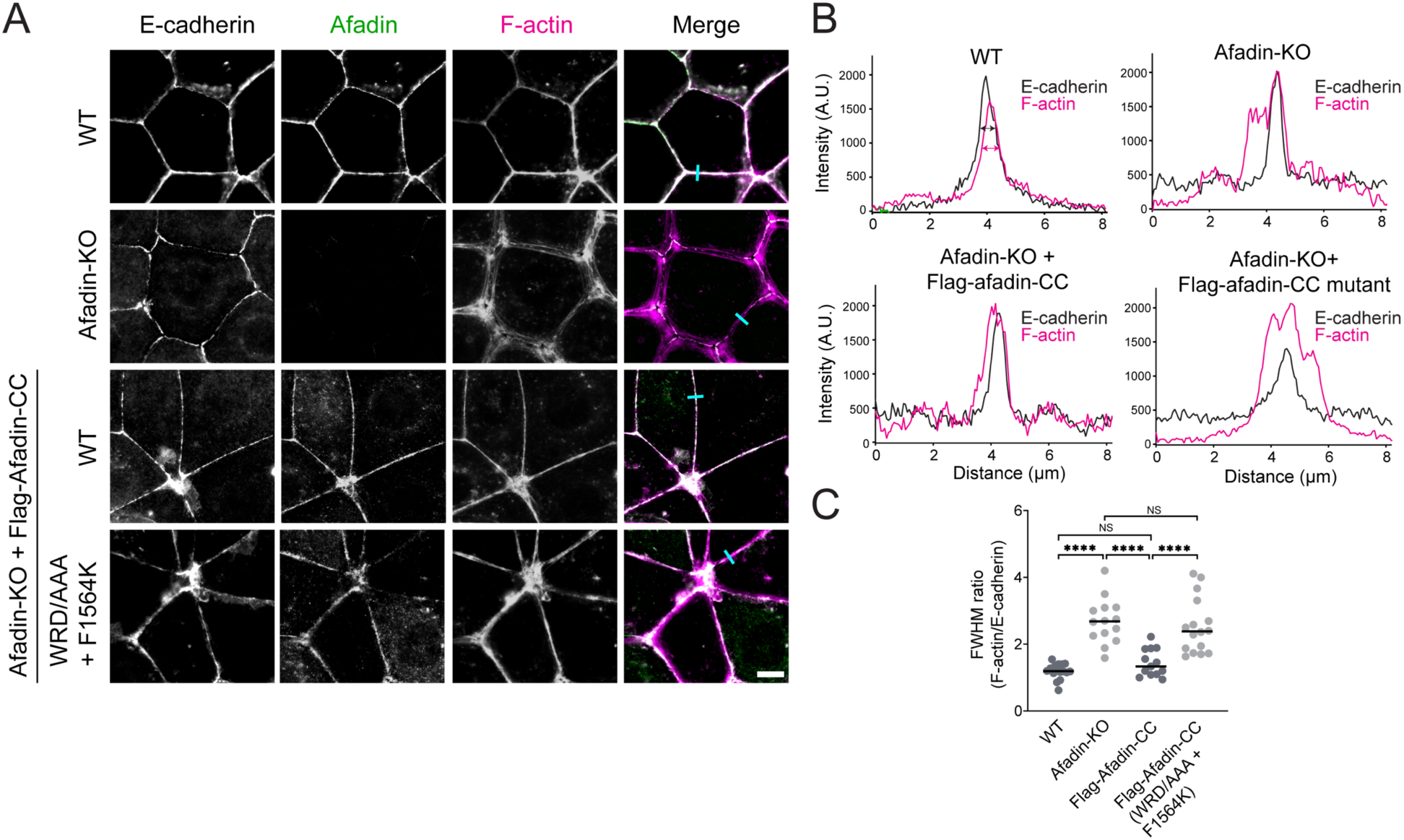
Afadin-CC is required for proper actomyosin organization at AJs. (A) Immunofluorescence staining of the indicated proteins at AJs in fixed EpH4 cells. Scale bar: 10 μm. (B) Line scan quantification of F-actin and E-cadherin signal along the indicated lines (cyan) in (A). Magenta and green double headed arrows indicate the full width at half maximum (FWHM) peak intensities for F-actin and E-cadherin, respectively. (C) Quantification of FWHM ratio of F-actin to E-cadherin. Bars represent means. Data represent three independent experiments (13 ≤ n ≤ 15) compared using one-way ANOVA with Fisher’s LSD test. NS, p ≥ 0.05; ****p < 0.0001.

### Afadin-CC promotes cooperative F-actin binding by the cadherin-catenin–vinculin supra-complex

Our finding that afadin-CC bridges adjacent αE-catenin ABDs along F-actin strands led us to hypothesize that it would promote cooperative binding of the tetrameric complex (Eβα(CA)-vinculin (T12-IV)) to F-actin, a behavior previously observed for the isolated αE-catenin ABD but reported to be absent for the trimeric Eβα complex^13,22^. We implemented a Total Internal Reflection Fluorescence (TIRF) microscopy assay to visualize the tetramer’s F-actin binding dynamics (Figure 6A). Adding a Halo tag to the N-terminus of E-cadherin for fluorescent labelling did not affect the tetramer’s weak intrinsic F-actin binding in co-sedimentation assays (Figure S2B). Correspondingly, we observed sparsely distributed binding events by TIRF, visible as small puncta of tetramer signal on individual filaments (Figure 6B). Addition of afadin-CC elicited a striking increase in tetramer binding, which occurred through the formation of discrete patches along filaments (Figure 6B). Time lapse imaging revealed bidirectional patch elongation, a hallmark of cooperative filament binding (Figure 6C). Fitting a saturation binding curve supports cooperative binding (K_d_ = 0.94 µM; Hill coefficient = 2.7) similar to that previously reported for the isolated αE-catenin ABD (Figure 6D)^13^. These data suggest that afadin-CC potentiates αE-catenin’s intrinsic cooperative F-actin binding in the context of higher-order adhesion assemblies, likely by both relieving autoinhibition through afadin-CC-N and by mediating direct contacts between F-actin bound ABDs through afadin-CC-C.

**Figure 6.**
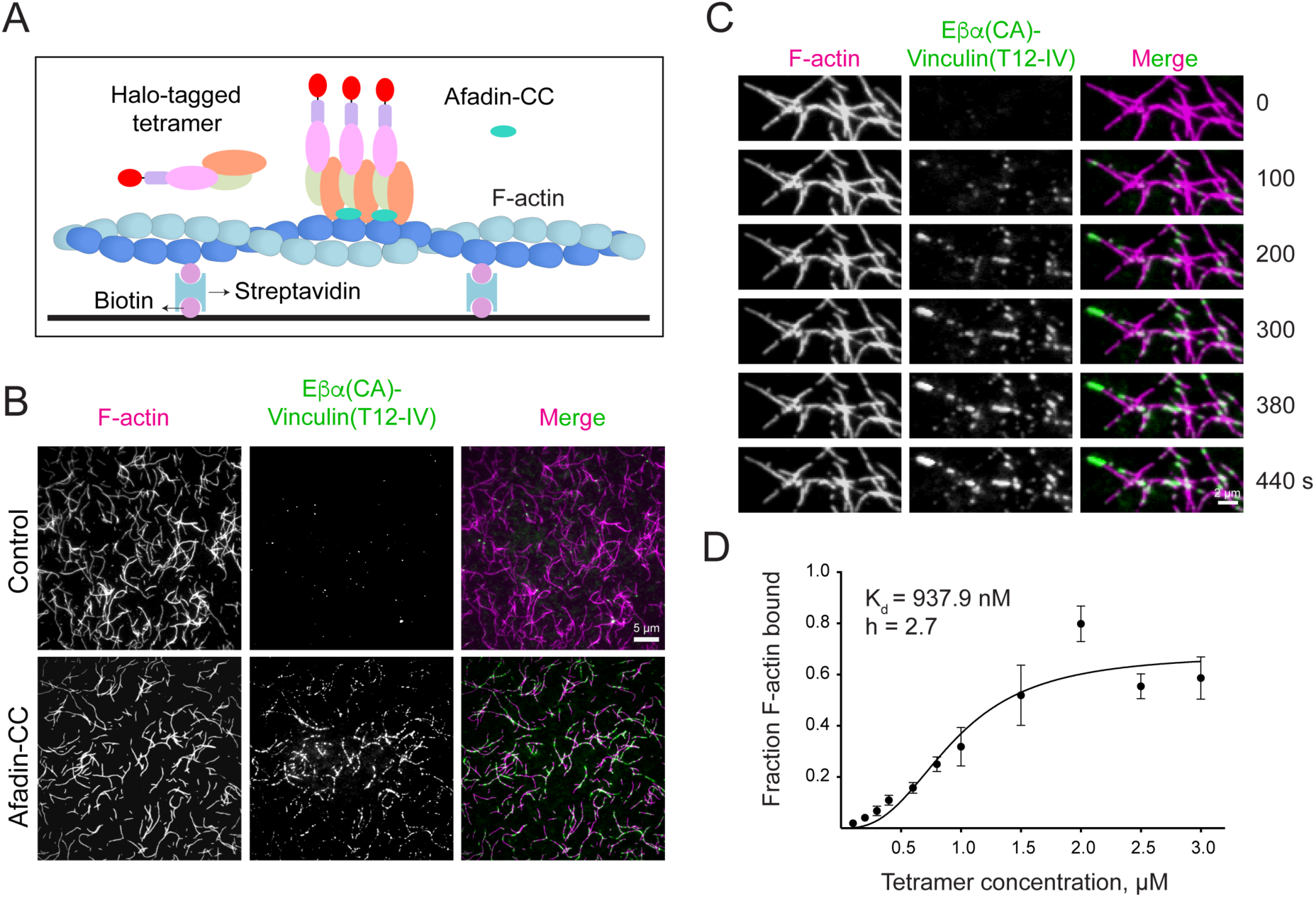
Afadin-CC stimulates cooperative binding of the tetrameric complex to F-actin. (A) Schematic of TIRF microscopy assay. (B) TIRF snapshots of Atto488 labeled actin (magenta) and Janelia Fluor 549 labeled Eβα(CA)-vinculin(T12-IV) (green) in the absence (top) or presence (bottom) of unlabeled afadin-CC. Scale bar, 5 μm. (C) Montage of tetramer patches in the presence of afadin-CC. Scale bar, 2 μm. (D) Fraction F-actin bound plotted against concentration of the tetramer in the presence of four-fold excess afadin-CC.

### Supra-complex binding along individual F-actin strands is associated with filament bending

In our previous studies of force-activated F-actin binding by the isolated αE-catenin ABD, we speculated that inter-ABD cooperative contacts could facilitate the protein’s discrimination of mechanical transitions in F-actin^23^. We recently experimentally confirmed this hypothesis in a study performed contemporaneously with the work presented here^53^. Our finding that afadin-CC enhances these contacts, as well as our recovery of 2D classes featuring curved F-actin decorated along a single strand from our cryo-EM dataset, led us to hypothesize the full pentamer could biochemically stabilize architectural remodeling of the filament. We therefore conducted detailed structural analysis of the segments contributing to curved F-actin classes, resulting in a 4.1 Å reconstruction (Figure 7A). Consistent with 2D analysis, the F-actin in this reconstruction features notable curvature, as well as density for the αE-catenin ABD and afadin-CC solely present along a single strand, occupying the convex side of the curve. Docking the atomic model of the pentamer-F-actin interface derived from the straight reconstruction into this density map did not reveal any notable changes in local binding contacts, suggesting that recognition and stabilization of deformed F-actin likely depends on the discrimination of longer-range architectural features by cooperative multi-pentamer assemblies.

**Figure 7.**
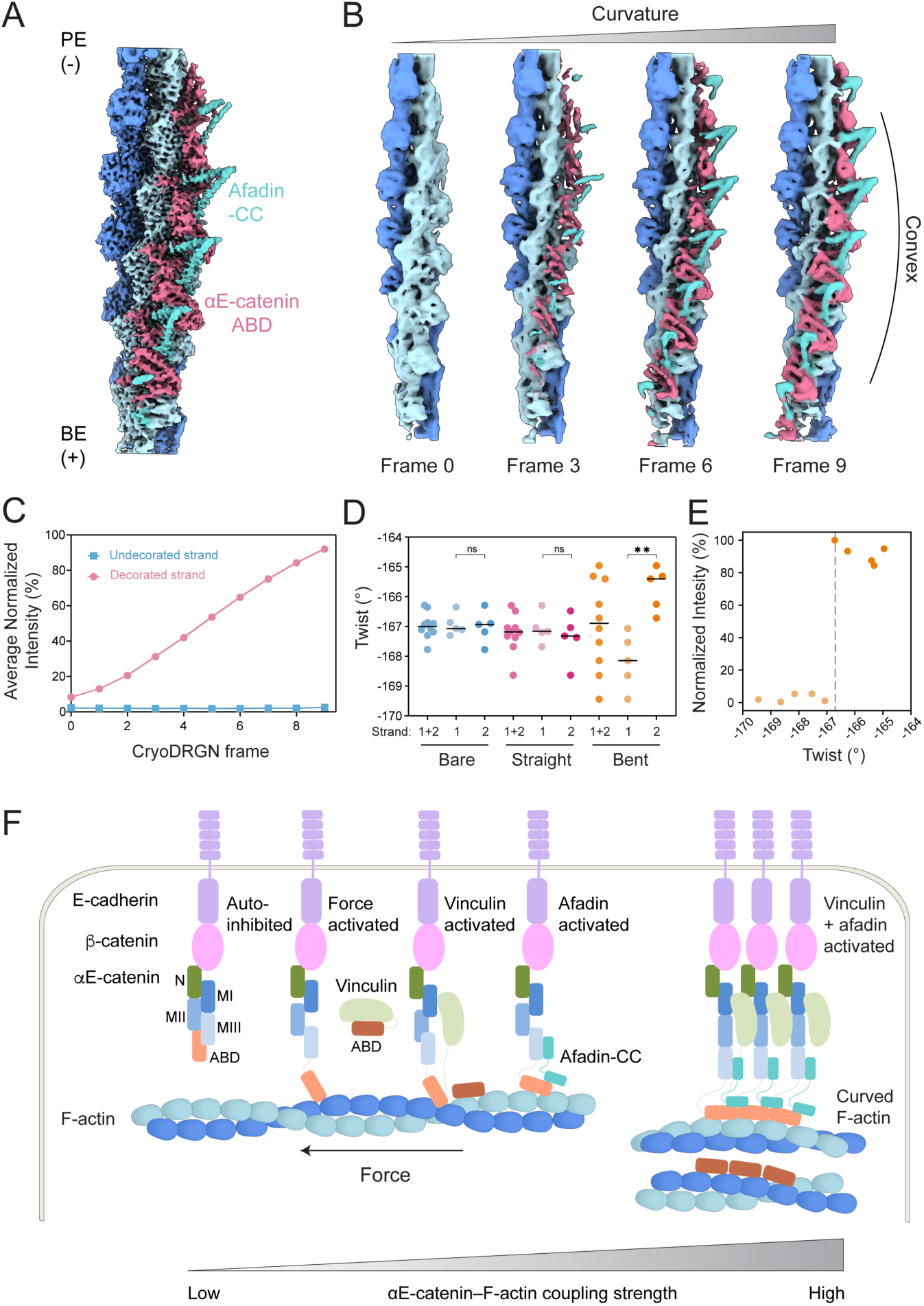
Cooperative binding of the pentameric complex is correlated with F-actin curvature. (A) 4.1 Å resolution density map that resolves αE-catenin ABDs and bridging afadin-CCs bound along a single F-actin strand. (B) Frames from cryoDRGN variability analysis capture coupling between increasing filament curvature and complex binding. (C) Quantification of average complex intensity along each F-actin strand across the cryoDRGN trajectory. (D) Quantification of the local twist of the bare filament (“bare”), straight filament with both strands decorated (“straight”), and the most curved filament (frame 9) from cryoDRGN analysis (“curved”). Circles represent the twist measured at each protomer index of the 10 central subunits. Strand 2 corresponds to the decorated strand of the curved filament. Bars represent means. Data were analyzed by one-way ANOVA with Tukey’s multiple comparison test: NS, p ≥ 0.05; **p < 0.01. (E) Quantification of complex intensity vs. local twist at each protomer index in the curved reconstruction from D. Vertical dashed line indicates twist of canonical straight F-actin. (F) Schematic model of the tunable coupling strength between the cadherin-catenin complex and F-actin, modulated by mechanical force, vinculin, and afadin.

To investigate the link between filament curvature and pentamer engagement, we performed variability analysis with cryoDRGN (Methods). Analysis of a volume series representing the first principal component of variation revealed a clear correlation between increasing curvature and the appearance of pentamer density along the convex strand, suggesting a link between filament deformation and cooperative binding along an individual F-actin strand (Figure 7B). To quantify this phenomenon, we measured the average normalized intensity of the pentamer density (a proxy for occupancy) along each strand for each frame (Figure 7C). To analyze architectural remodeling of the filament lattice accompanying this deformation, we also measured the instantaneous helical twist and rise at each subunit index in the cryoDRGN frame featuring maximal curvature (frame 9). As points of comparison, from the same dataset we also asymmetrically reconstructed a bare filament region and a fully decorated straight filament region spanning an equivalent number of subunits (Figures S7A and S7B). As anticipated, twist is highly uniform for both the bare filament and the fully decorated straight filament. In contrast, although the overall average twist of the maximal curvature frame is similar to the straight filaments, the average twist of each strand is distinct in the maximal curvature volume, consistent with prior reports of twist-bend coupling in F-actin^54^. The highly decorated strand is substantially under-twisted (Figure 7D), leading to a correlation between subunits featuring a lower twist magnitude and high pentamer intensity (Figure 7E). Collectively, this suggests cooperative pentamer binding either induces or stabilizes filament curvature, with preferential binding occurring along the convex face of curved filament regions.

In our contemporary study of the isolated αE-catenin ABD engaging F-actin in the presence of myosin-generated forces, we find that the ABD preferentially engages positions featuring extended rise^53^. To assess whether the pentameric complex induces or stabilizes similar lattice rearrangements, we analyzed the correlation between instantaneous helical rise and pentamer intensity. While the bare filament displayed uniform helical rise, both the fully decorated straight filament and the maximal curvature filament displayed increased rise along one strand and decreased rise along the other (Figure S7C), nevertheless maintaining an overall average similar to bare F-actin^54^. In the curved filament, the subunits featuring increased rise are correlated with high pentamer intensity (Figure S7D), consistent with the behavior of the isolated ABD in the presence of myosin forces^53^. However, in the fully straight filament, all subunits are equivalently decorated, suggesting rise changes are not the sole determinant of asymmetric engagement by the pentamer. One potential explanation for this phenomenon is that the fully decorated reconstruction represents a mixture of different species featuring variable pentamer occupancy which we were unable to sort via current 3D classification or variability methods. Nevertheless, these data are broadly consistent with afadin-CC stabilizing cooperative, asymmetric binding by the cadherin-catenin complex to curved F-actin, reminiscent of force-activated binding by the αE-catenin ABD.

## Discussion

This study provides biochemical and structural insights into how the intrinsic actin-binding activity of the core cadherin-catenin complex can be modulated by additional adhesion factors, revealing mechanisms which could tune AJ-cytoskeleton coupling strength for function across varying mechanical conditions. Our biochemical data are broadly consistent with a model in which the association of vinculin and afadin-CC cumulatively relieve autoinhibition of αE-catenin’s ABD, with the convergence of both proteins supporting substantially higher F-actin engagement than either factor alone. We further found evidence of supra-complex composition-dependent interplay between the vinculin ABD and the αE-catenin ABD. The vinculin ABD serves as the dominant actin-binding moiety in the tetrameric cadherin-catenin–vinculin complex, consistent with a recent study^47^, while the αE-catenin ABD plays a predominant role in the pentameric cadherin-catenin–vinculin–afadin complex. This suggests the vinculin and αE-catenin ABDs may have dynamically varying functions as AJs undergo compositional and mechanical remodeling *in vivo,* an important topic for future mechanistic cell biology studies, which can be guided by the biophysical framework introduced here.

Our structural studies furthermore show direct modulation of the αE-catenin–F-actin interface by the afadin-CC, providing an additional mechanism for stabilizing the cadherin-catenin complex’s F-actin binding. Afadin-CC engages flexible αE-catenin ABD elements implicated in force-based enhancement of αE-catenin’s F-actin binding^23^, supporting a model in which afadin biochemically coopts α-catenin’s mechanical regulatory mechanisms to achieve constitutively strong, stable binding. Interpreted through the framework where H1 undocking from the αE-catenin ABD helical bundle mediates catch-bonding, while CTE dynamics mediate force-activated binding, our data predict that force is unlikely to additionally stabilize F-actin binding once the pentameric complex featuring afadin-CC has fully engaged. Nevertheless, by pre-organizing these structural elements in the αE-catenin ABD, force is likely to promote afadin association with the complex. Functionally, this could facilitate persistent maximal AJ-cytoskeletal coupling at cell-cell junctions bearing consistently high mechanical loads. Selective disruption of specific afadin-CC–αE-catenin ABD contacts we have identified, which are highly conserved, can facilitate precise dissection of the functions of this interface in tissue dynamics across species.

Additionally, we find that afadin-CC mediates contacts between neighboring αE-catenin ABDs, promoting the formation of cooperative assemblies along individual F-actin strands in filament segments featuring nanoscale curvature. While to our knowledge this binding mode has not previously been described, other actin-associated proteins, including IQGAP and septins, have been reported to promote F-actin bending, albeit at the micron rather than nanometer scale^55,56^. Structural studies of these factors in complex with F-actin will be required to decipher whether they employ similar mechanisms. While we speculate that the pentamer’s recognition / stabilization of F-actin curvature represents further cooption of mechanical regulation by afadin, it could also feasibly serve additional functions. In cells, the complex is localized to the plasma membrane, where individual strands of actin filaments adjacent to the membrane are locally accessible^21,36,57^. Forming assemblies along F-actin strands thus represents a cooperativity mechanism compatible with physiological geometric restrictions. Moreover, afadin is frequently localized to sites featuring high local membrane curvature, such as tricellular junctions^37,58^, where stabilizing F-actin curvature could conceivably enhance AJ–cytoskeleton linkages by aligning the contour of filaments with that of the membrane. Nanoscale imaging studies, such as cryo-electron tomography of AJs with varying architectures and afadin abundance, will be necessary to dissect the detailed functional links between local F-actin morphology and AJ-cytoskeleton coupling.

## Acknowledgements

We gratefully acknowledge Johanna Sotiris, Honkit Ng, and Mark Ebrahim at the Rockefeller University (RU) Evelyn Gruss Lipper Cryo-EM Resource Center (CEMRC) for their support in data collection, and Ariel Pan at RU for performing F-actin purification. We thank Bob Weinberg for providing the cDNA of mouse E-cadherin (plasmid #18804) through Addgene, as well as Valeri Vasioukhin for providing the cDNAs of mouse β-catenin (plasmid #20140) and αE-catenin (plasmid #20139). We acknowledge Professor Yoshimi Takai (Kobe University, Japan) for generously supplying wild-type and afadin knockout EpH4 cell lines. We are grateful to Professor Kun-Liang Guan (University of California, San Diego) for providing HEK293T cells and the Dox-inducible pCDH vectors for lentiviral production. This work was funded by grants from the NIH (R01GM141044), the Alfred P. Sloan Foundation (G-2020-14047), and the Stavros Niarchos Foundation Institute for Global Infectious Disease Research at the Rockefeller University to G.M.A.

## Resource Availability

All resources and reagents reported in this study are freely available, and requests should be directed to the lead contact, Gregory M. Alushin (galushin@.rockefeller.edu).

## Data Availability

The cryo-EM density maps, atomic model, and tomogram generated in this study have been deposited in the Electron Microscopy Data Bank (EMDB) and Protein Data Bank (PDB): F-actin binding interface of ⍺E-catenin ABD (cadherin-catenin complex) and afadin (EMD-47194, PDB:9DVA); ⍺E-catenin ABD (cadherin-catenin complex) and afadin bound to bent F-actin (EMD-47195); bare F-actin (EMD-47196); ⍺E-catenin ABD (cadherin-catenin complex) and afadin bound to straight F-actin (EMD-47197); tomogram of the cadherin-catenin–vinculin-afadin complex bound to F-actin (EMD-47198). The trained neural networks for denoising and semantically segmenting micrographs, atomic models and volumes used to generate synthetic datasets for network training, and cryoDRGN frames / docking models used for curvature vs. occupancy analysis are available at Zenodo: https://doi.org/10.5281/zenodo.13899747. All other data are presented in the manuscript.

## Code Availability

Custom code is available at https://www.github.com/alushinlab/afadin as open source.

## Author Contributions

R.G. and G.M.A. designed research. R.G. purified proteins, collected cryo-EM and cryo-ET datasets, and performed biochemical, cell biology, and TIRF studies. M.J.R. and R.G. processed cryo-EM data. R.G. built atomic models. R.G. reconstructed the tomograms. X.S. performed Hill equation fitting analysis. R.G. and G.M.A. wrote the paper with input from all other authors.

---

**Figure S1.**
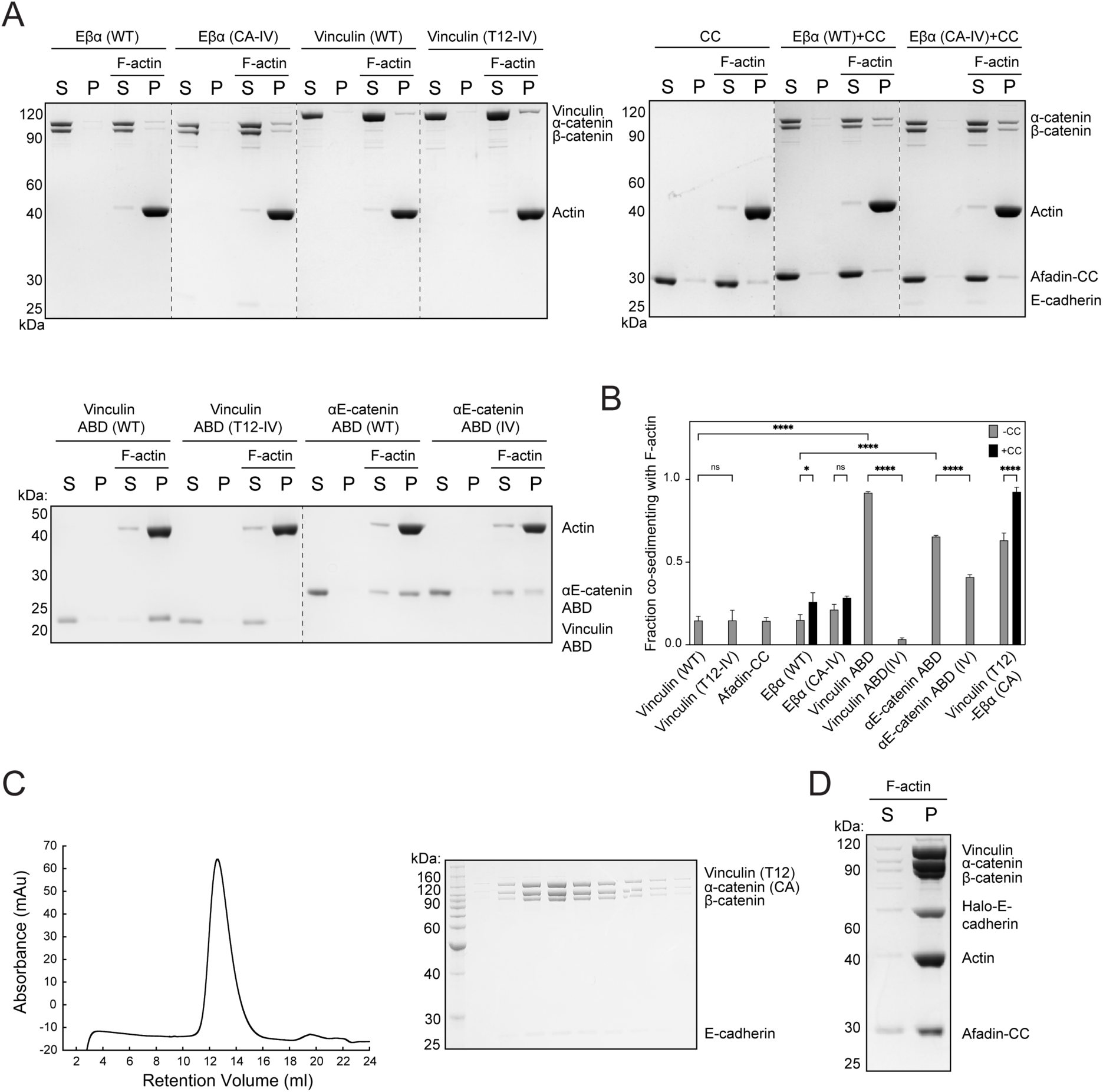
F-actin binding activities of the cadherin-catenin complex, vinculin and afadin-CC. (A) Co-sedimentation assays of indicated proteins and protein complexes with F-actin. Vinculin(WT), wild-type vinculin; α(WT), wild-type αE-catenin. All other abbreviations are the same as in Figure 1. Dotted lines indicate stitching interfaces between gels. (B) Quantification of A. Data are presented as mean ± SD of three independent experiments, compared via two-way ANOVA with Tukey’s multiple comparison test. NS, p ≥ 0.05; *p < 0.05; ****p < 0.0001. (C) Left: Size exclusion chromatography of the tetrameric vinculin(T12)-Eβα(CA) complex. Right: analysis of peak fractions by SDS-PAGE. (D) Co-sedimentation of Halo-tagged Eβα(CA)-vinculin(T12-IV) with F-actin in the presence of afadin-CC.

**Figure S2.**
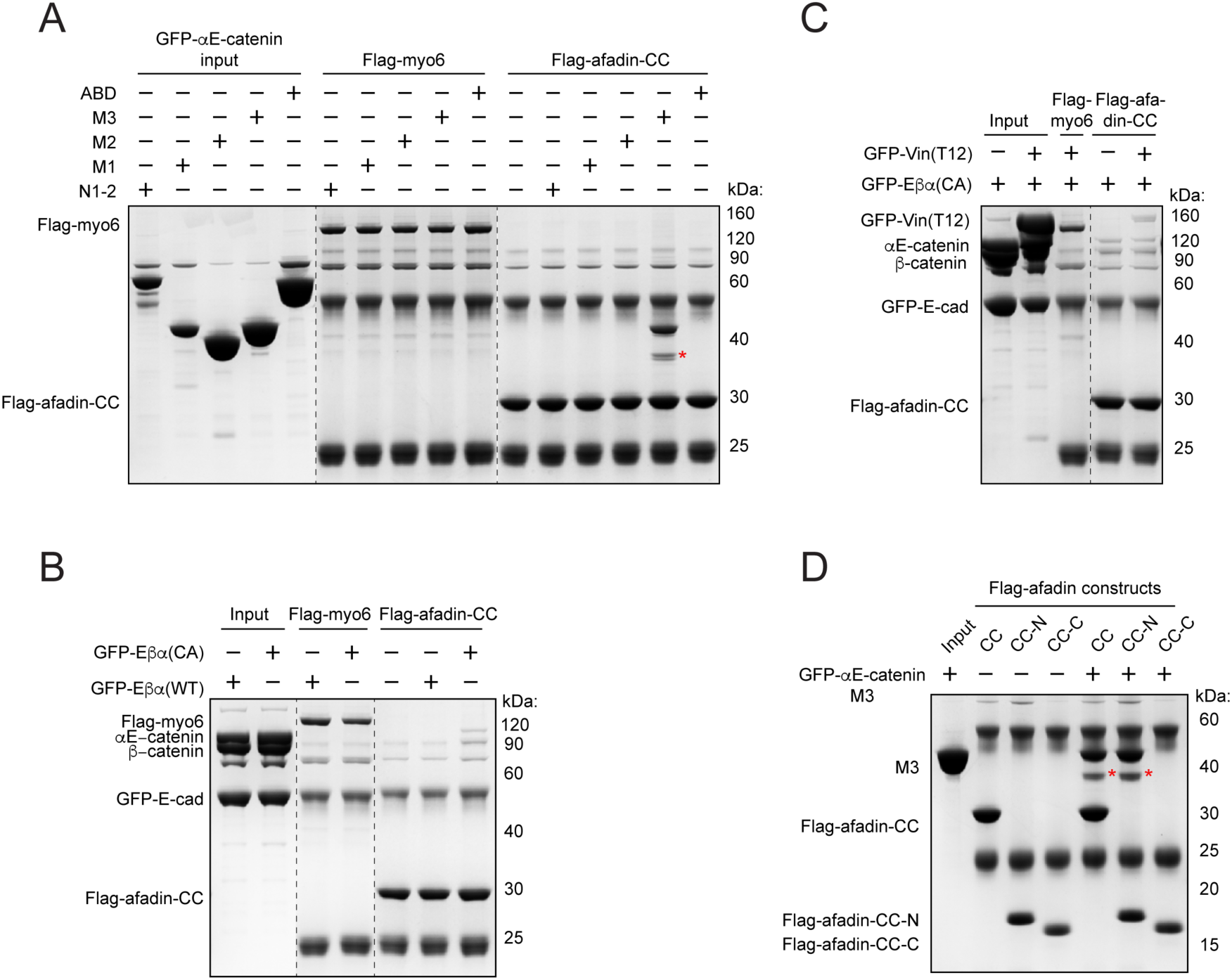
Afadin-CC forms a pentameric complex with vinculin (T12)-Eβα(CA) through its direct interaction with the M3 domain of αE-catenin. (A) Pull-down assay demonstrates specific binding between afadin-CC and the M3 domain of αE-catenin. The Flag-tagged motor domain of myosin-6 (Flag-myo6) was used as a negative control. The red asterisk indicates nonspecific binding. (B) Pull-down assay showing binding between Flag-tagged afadin-CC and GFP-tagged Eβα(CA). (C) Pull-down assay showing the interaction between afadin-CC and Eβα(CA)-vinculin(T12). (D) Pull-down assay showing the association between afadin-CC-N and the M3 domain of αE-catenin. Red asterisks indicate nonspecific binding.

**Figure S3.**
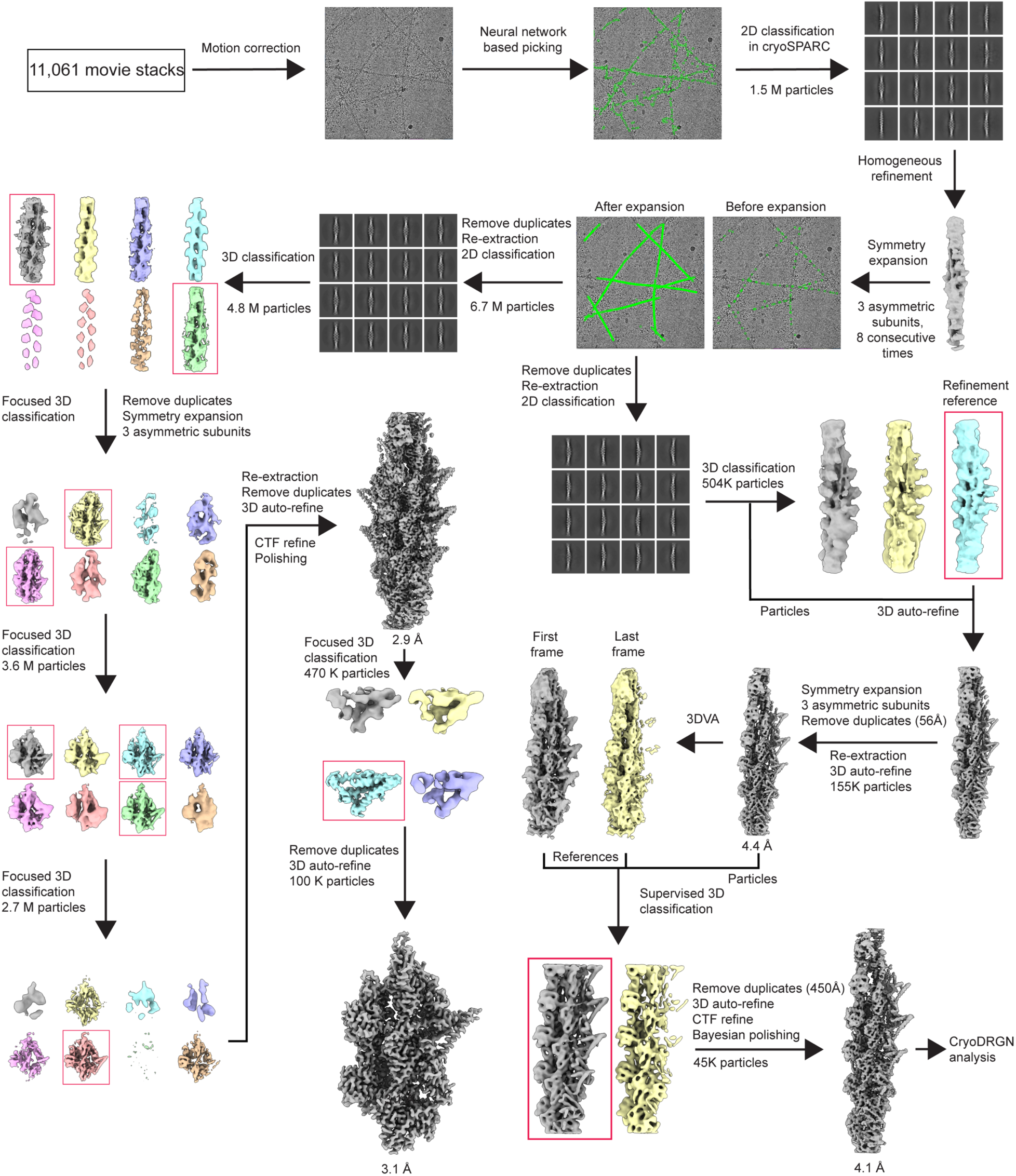
Cryo-EM data processing workflow. Scheme outlining the combined neural-network based picker and symmetry expansion approach used to capture heterogeneously decorated filaments in the cryo-EM images, followed by single particle processing and variability analysis

**Figure S4.**
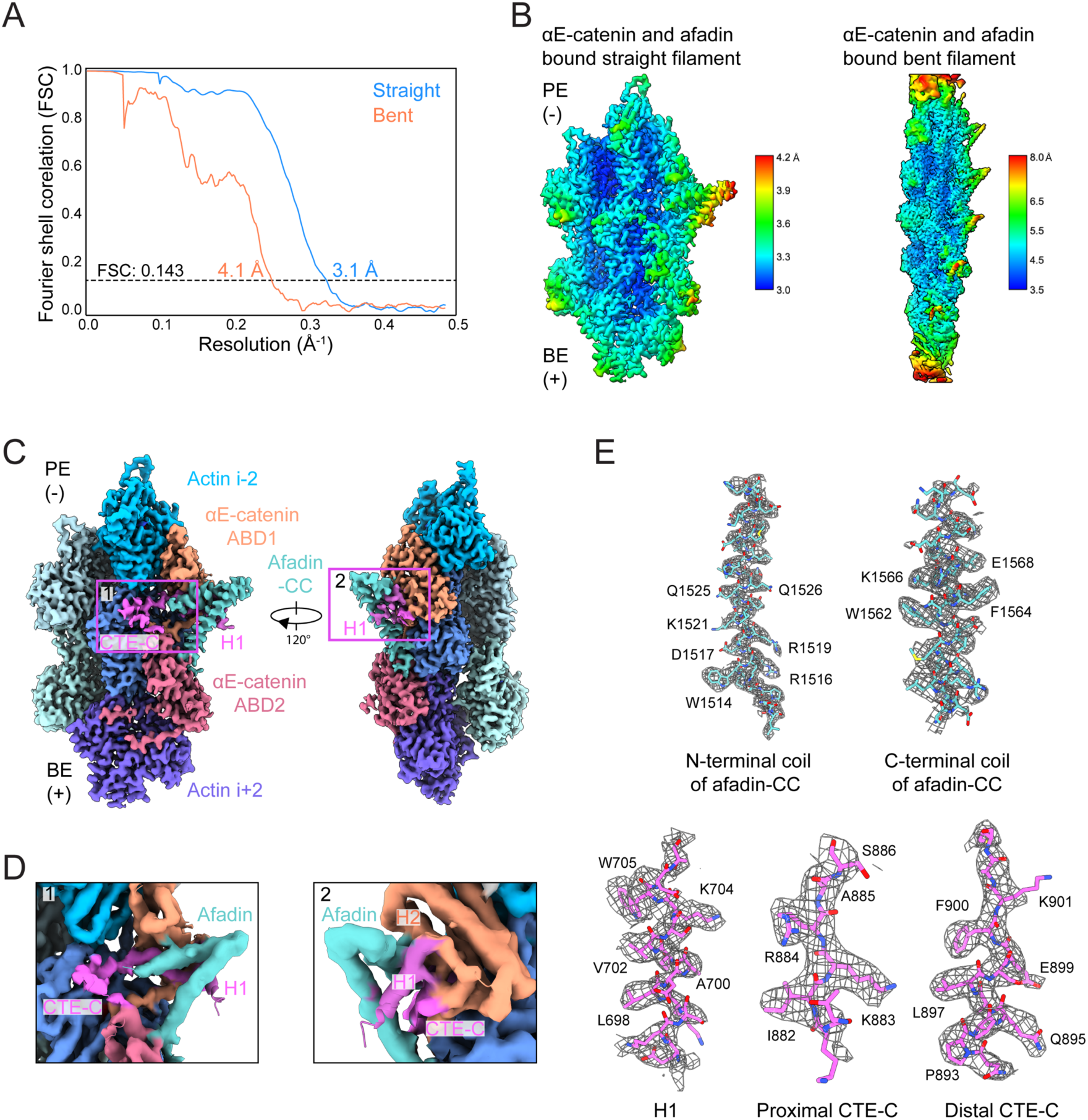
Resolution assessment and density features of cryo-EM reconstructions. (A) Gold-standard Fourier Shell Correlation (FSC) curves for the straight and curved pentamer-bound F-actin reconstructions. (B) Local resolution estimation of the two reconstructions. (C) Views of the straight pentamer-bound F-actin cryo-EM map, highlighting densities corresponding to flexible αE-catenin ABD segments stabilized by afadin-CC. (D) Detail views of boxed regions in C. The map was low-pass filtered to 6 Å to facilitate visualization. (E) Segmented cryo-EM map densities and atomic models of the afadin-CC and the αE-catenin structural elements it stabilizes.

**Figure S5.**
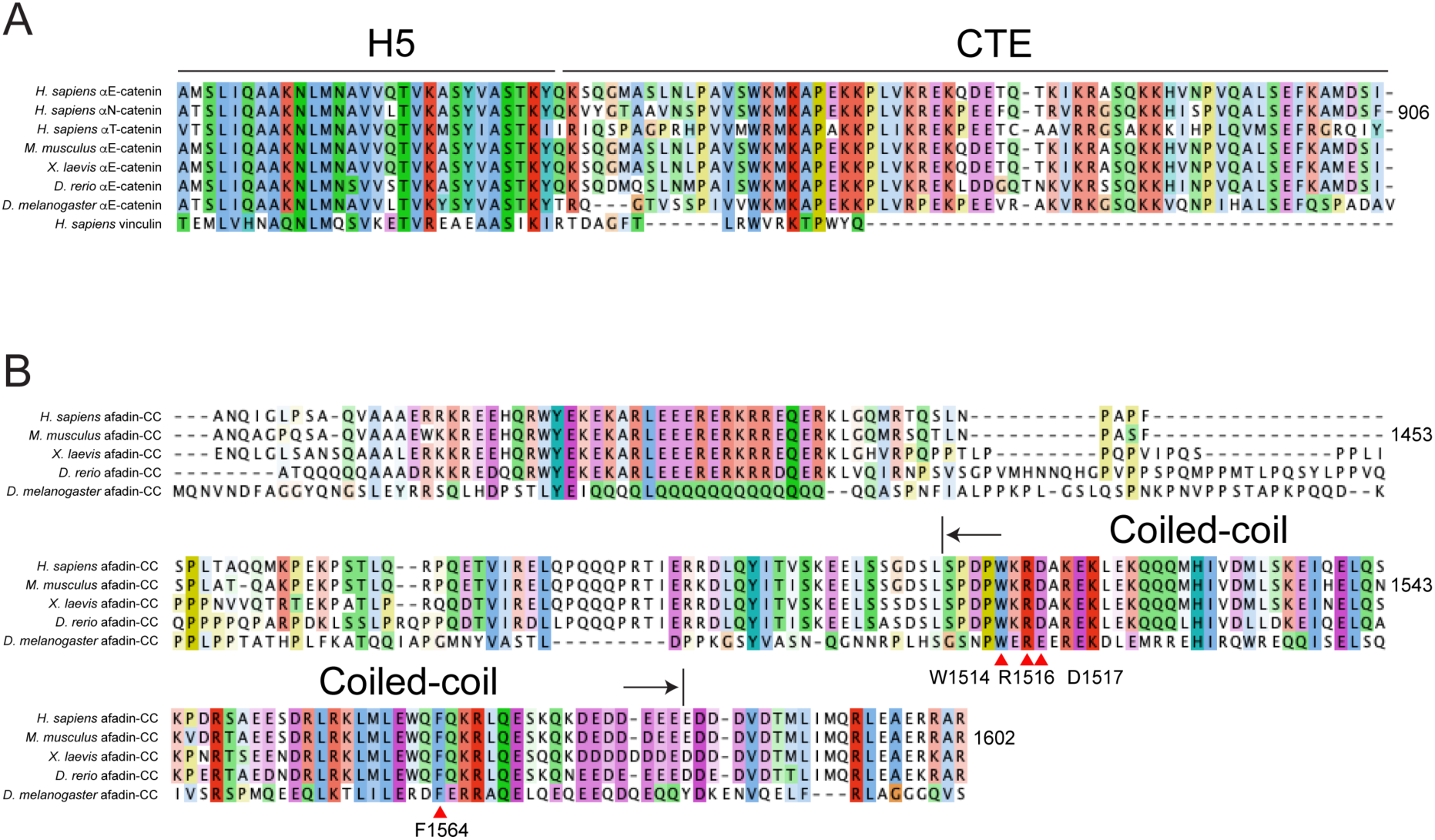
Sequence alignments of the α-catenin CTE and afadin-CC. (A) Sequence alignment of the α-catenin CTE among different species and isoforms. Sequences used for alignment are from *H. sapiens* αE-catenin (NP_001310911.1), *H. sapiens* αN-catenin (NP_004380.2), *H. sapiens* αT-catenin (NP_001120856.1), *M. musculus* αE-catenin (NP_033948.1), *X. laevis* αE-catenin (NP_001084100.1), *D. rerio* αE-catenin (NP_571531.1), *D. melanogaster* αE-catenin (NP_524219.1) and *H. sapiens* vinculin (NP_003364.1). (B) Sequence alignment of afadin’s coiled-coil region. Aligned sequences are from *H. sapiens* afadin (NP_001353249.1), *M. musculus* afadin (NP_034936.1), *X. laevis* afadin (NP_001171575.1), *D. rerio* afadin (XP_021324351.1), and *D. melanogaster* canoe (NP_524232.2). Both alignments are colored by sequence conservation. Alignments were performed using Clustal Omega and visualized with Jalview.

**Figure S6.**
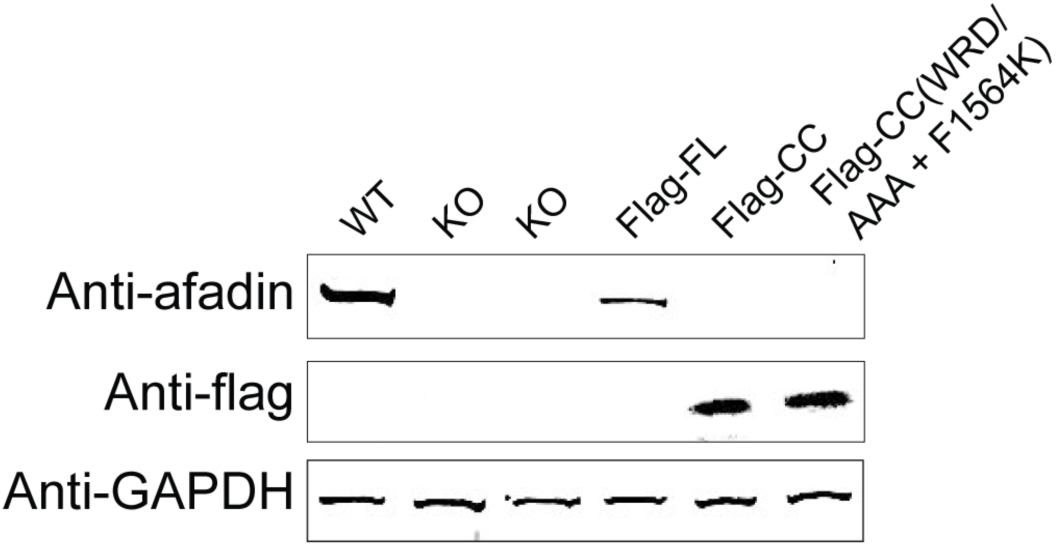
Western blots showing afadin knockout and overexpression efficiency. Experiments were performed in EpH4 cells.

**Figure S7.**
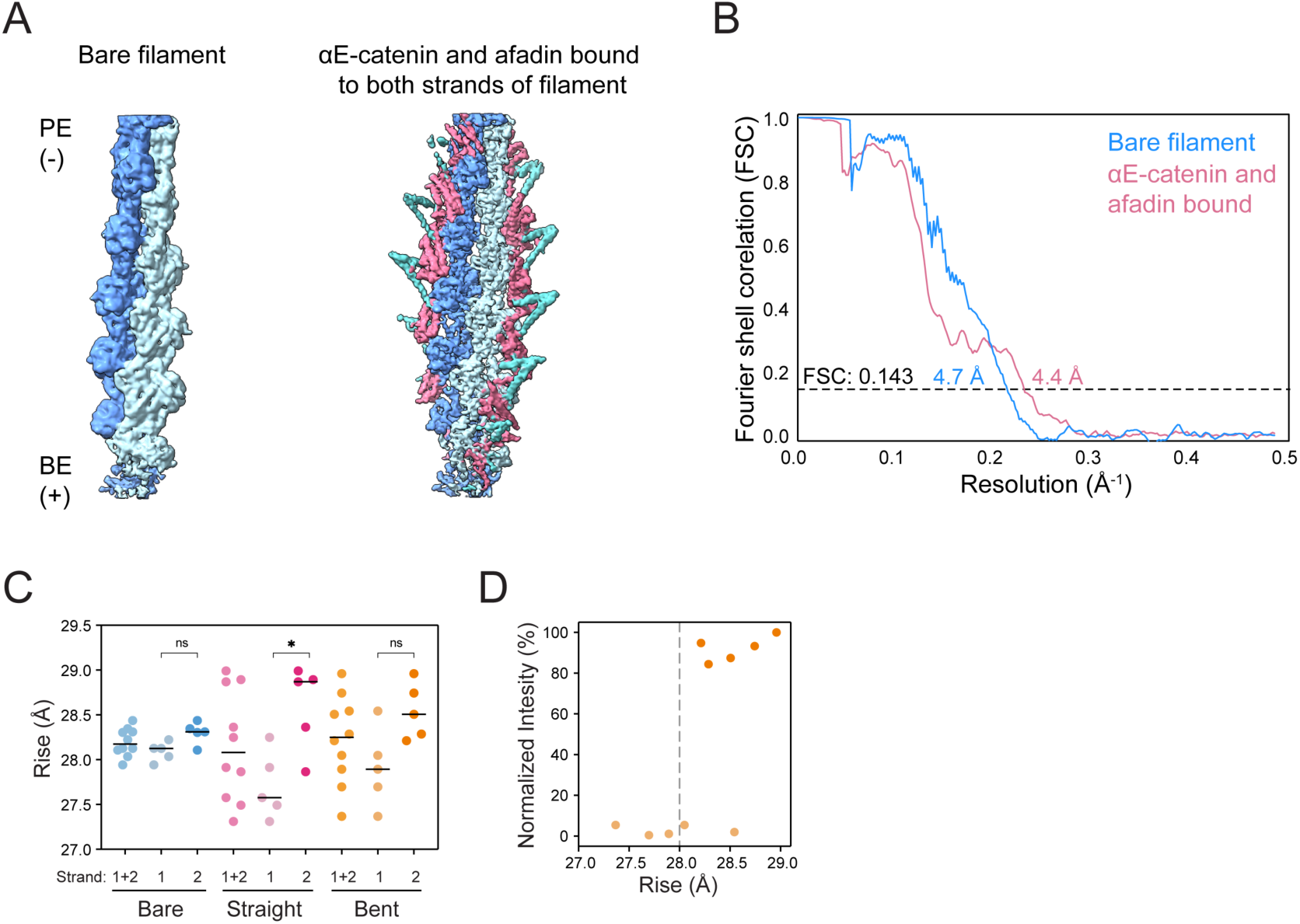
Cryo-EM structures of bare F-actin and pentamer bound to both strands of F-actin spanning 13 actin subunits. (A) 4.7 Å and 4.4 Å resolution cryo-EM density maps of bare F-actin (left) and pentamer bound to both strands of F-actin (right). (B) FSC curves for the bare F-actin and pentamer bound to both strands of F-actin reconstructions. (C) Quantification of the local rise of the bare filament (“bare”), straight filament with both strands decorated (“straight”), and the most curved filament (frame 9) from cryoDRGN analysis (“curved”). Circles represent the rise measured at each protomer index of the 10 central subunits. Strand 2 corresponds to the decorated strand of the curved filament. Bars represent means. Data were compared by one-way ANOVA with Tukey’s multiple comparison test: NS, p ≥ 0.05; *p < 0.05. (D) Quantification of complex intensity vs. local rise at each protomer index in the curved reconstruction from C. Vertical dashed line indicates rise of canonical straight F-actin.

**Table S1.**
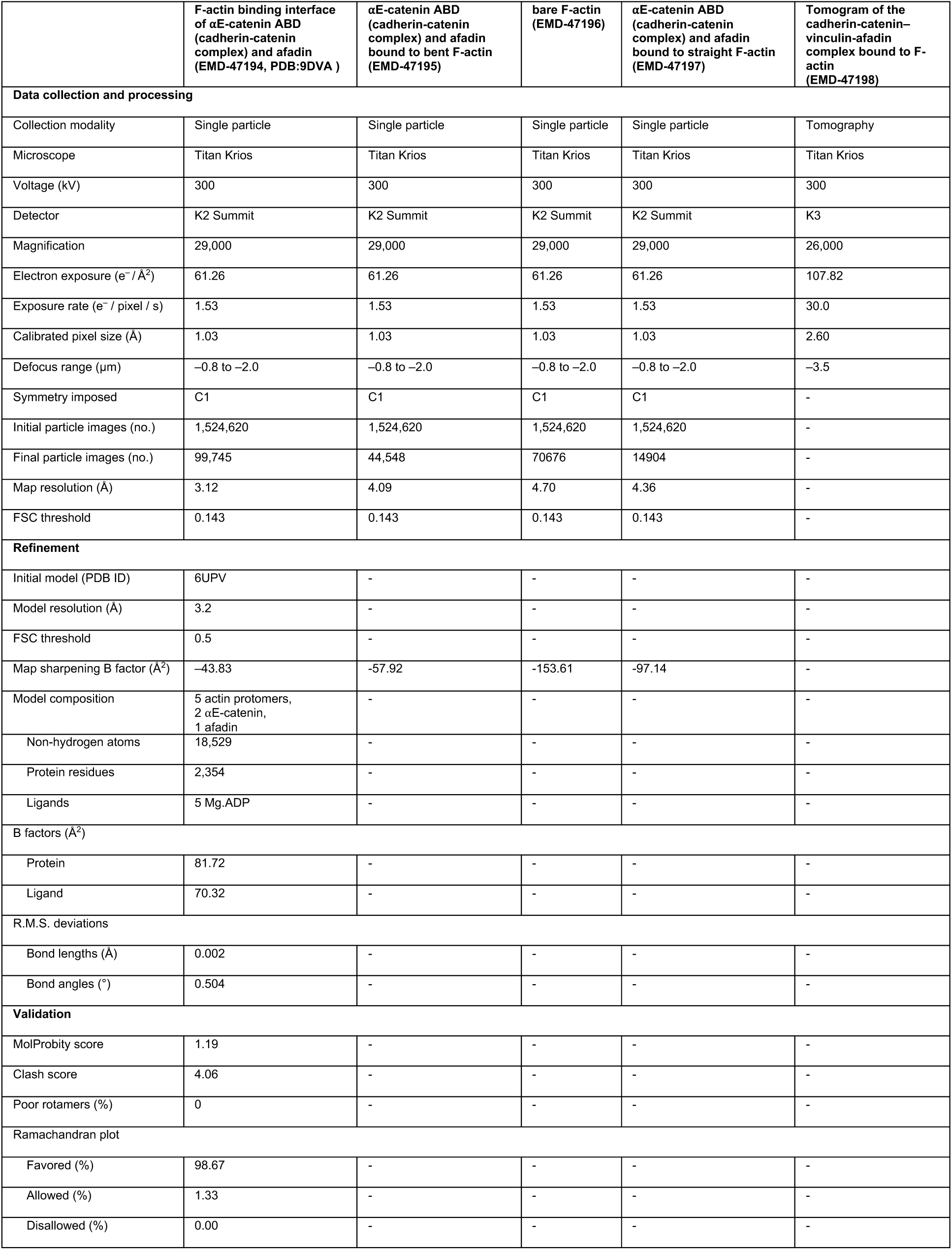
Cryo-EM data collection, refinement and validation statistics.

| | F-actin binding interface of αE-catenin ABD (cadherin-catenin complex) and afadin (EMD-47194, PDB:9DVA ) | αE-catenin ABD (cadherin-catenin complex) and afadin bound to bent F-actin (EMD-47195) | bare F-actin (EMD-47196) | αE-catenin ABD (cadherin-catenin complex) and afadin bound to straight F-actin (EMD-47197) | Tomogram of the cadherin-catenin-vinculin-afadin complex bound to F-actin (EMD-47198) |
| --- | --- | --- | --- | --- | --- |
| **Data collection and processing** |  |  |  |  |  |
| Collection modality | Single particle | Single particle | Single particle | Single particle | Tomography |
| Microscope | Titan Krios | Titan Krios | Titan Krios | Titan Krios | Titan Krios |
| Voltage (kV) | 300 | 300 | 300 | 300 | 300 |
| Detector | K2 Summit | K2 Summit | K2 Summit | K2 Summit | K3 |
| Magnification | 29,000 | 29,000 | 29,000 | 29,000 | 26,000 |
| Electron exposure ($e^- / \text{\AA}^2$) | 61.26 | 61.26 | 61.26 | 61.26 | 107.82 |
| Exposure rate ($e^- / \text{pixel} / \text{s}$) | 1.53 | 1.53 | 1.53 | 1.53 | 30.0 |
| Calibrated pixel size ($\text{\AA}$) | 1.03 | 1.03 | 1.03 | 1.03 | 2.60 |
| Defocus range ($\mu\text{m}$) | -0.8 to -2.0 | -0.8 to -2.0 | -0.8 to -2.0 | -0.8 to -2.0 | -3.5 |
| Symmetry imposed | C1 | C1 | C1 | C1 | - |
| Initial particle images (no.) | 1,524,620 | 1,524,620 | 1,524,620 | 1,524,620 | - |
| Final particle images (no.) | 99,745 | 44,548 | 70676 | 14904 | - |
| Map resolution ($\text{\AA}$) | 3.12 | 4.09 | 4.70 | 4.36 | - |
| FSC threshold | 0.143 | 0.143 | 0.143 | 0.143 | - |
| **Refinement** |  |  |  |  |  |
| Initial model (PDB ID) | 6UPV | - | - | - | - |
| Model resolution ($\text{\AA}$) | 3.2 | - | - | - | - |
| FSC threshold | 0.5 | - | - | - | - |
| Map sharpening B factor ($\text{\AA}^2$) | -43.83 | -57.92 | -153.61 | -97.14 | - |
| Model composition | 5 actin protomers,<br>2 αE-catenin,<br>1 afadin | - | - | - | - |
| Non-hydrogen atoms | 18,529 | - | - | - | - |
| Protein residues | 2,354 | - | - | - | - |
| Ligands | 5 Mg,ADP | - | - | - | - |
| **B factors ($\text{\AA}^2$)** |  |  |  |  |  |
| Protein | 81.72 | - | - | - | - |
| Ligand | 70.32 | - | - | - | - |
| **R.M.S. deviations** |  |  |  |  |  |
| Bond lengths ($\text{\AA}$) | 0.002 | - | - | - | - |
| Bond angles ($^\circ$) | 0.504 | - | - | - | - |
| **Validation** |  |  |  |  |  |
| MolProbity score | 1.19 | - | - | - | - |
| Clash score | 4.06 | - | - | - | - |
| Poor rotamers (%) | 0 | - | - | - | - |
| **Ramachandran plot** |  |  |  |  |  |
| Favored (%) | 98.67 | - | - | - | - |
| Allowed (%) | 1.33 | - | - | - | - |
| Disallowed (%) | 0.00 | - | - | - | - |

**Table S2:**
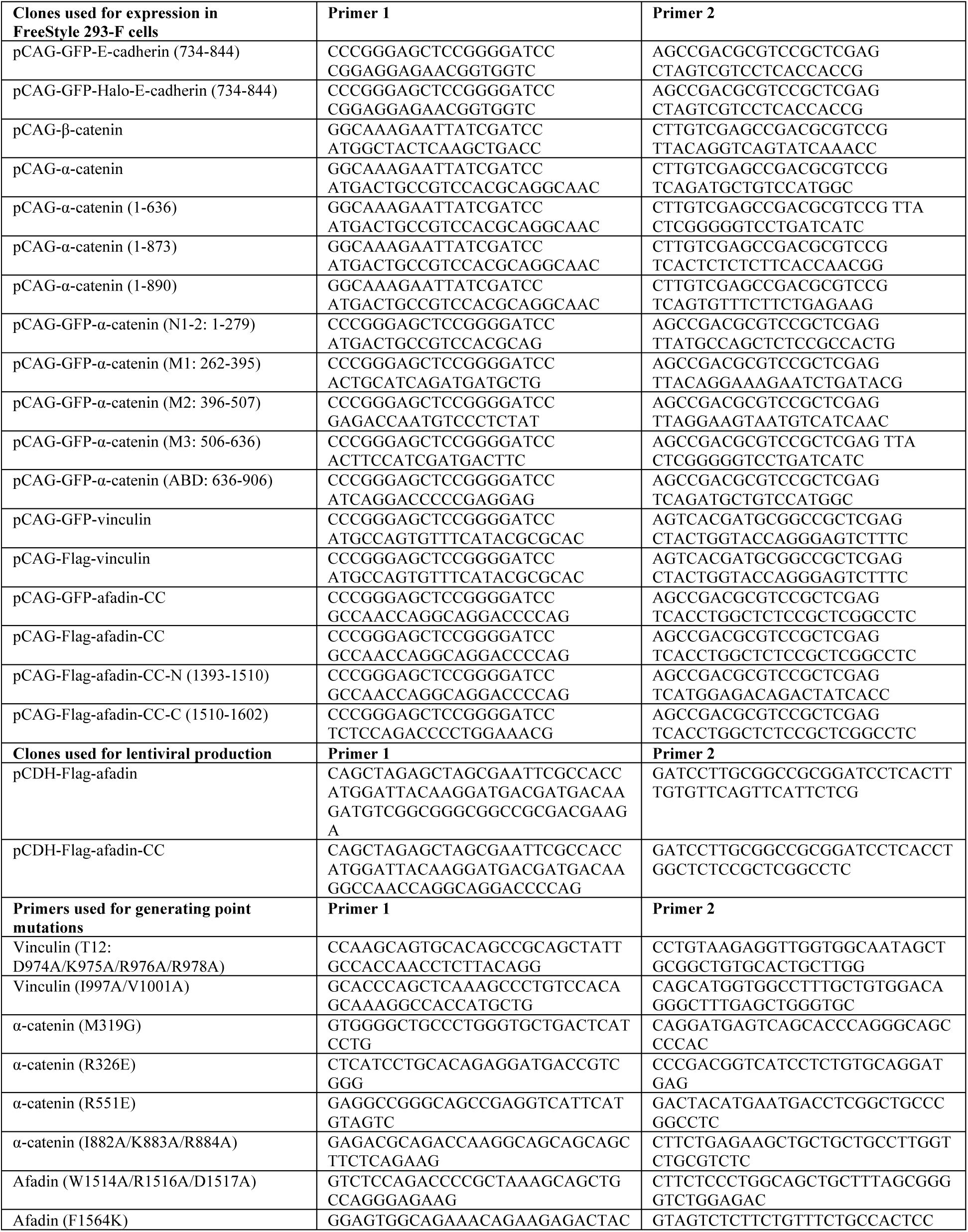
Constructs and Primers.

## Supplementary Video Legends

**Video S1. Tomogram of the pentameric supra-complex binding along actin filaments**.

**Video S2. Morph of the αE-catenin ABD model between the pre-bound and post-bound states.** The αE-catenin ABD model in the pre-bound state is from the AlphaFold2 predicted structure of the full length mouse αE-catenin (AF-P26231-F1), which contains the entire CTE-C.

**Video S3. Morph of the αE-catenin ABD model between the pre-bound and the afadin stabilized states.** The same pre-bound model was used as in Video S2.

## Methods

Unless otherwise noted, reagents were purchased from Sigma-Aldrich.

## Cell culture

Wildtype and afadin knockout EpH4 cell lines were a kind gift from Yoshimi Takai at Kobe University, Japan. HEK293T cells were generously provided by Kun-Liang Guan at the University of California, San Diego. The cells were maintained in DMEM medium (Thermo Fisher, 11995-065) supplemented with 10% Heat Inactivated Fetal Bovine Serum (Sigma, F4135) and 1x Antibiotic-Antimycotic (Thermo Fisher, 15240-062). FreeStyle 293-F cells (Thermo Fisher, R79007) were cultured in FreeStyle 293 expression medium (ThermoFisher, 12338018).

## Plasmids and cloning

The cDNAs of mouse E-cadherin (NM_004360.5, Addgene: #18804, gift of Bob Weinberg), β-catenin (NM_007614.3, Addgene #20140, gift of Valeri Vasioukhin), and αE-catenin (NM_009818.1, Addgene #20139, gift of Valeri Vasioukhin) were obtained from Addgene. The cDNA of mouse afadin (NM_010806.1) was purchased from Horizon Discovery (MMM1013-202707312). The cDNA of human vinculin (NM_003373.4) was PCR amplified from a cDNA library prepared from HEK 293T cells. Full-length coding sequences or fragments of E-cadherin, β-catenin, αE-catenin, vinculin, and afadin (Table S2) were individually cloned into a modified pCAG mammalian expression vector, either untagged or tagged with an N-terminal GFP or Flag tag (DYKDDDDK) followed by a TEV protease cleavage site^1^. To express GFP and Halo-tagged E-cadherin, a Halo tag was inserted immediately after the TEV cleavage site. For lentiviral production, the cDNAs of afadin constructs with an N-terminal Flag tag were cloned into a doxycycline-inducible pCDH-EF1a-MCS vector carrying a puromycin selection marker. The reverse tetracycline-responsive transcriptional activator (rtTA) was expressed in a separate pCDH vector with G418 selection. Both vectors were generously provided by Kun-Liang Guan at the University of California, San Diego. All wild-type and mutant clones were generated by Gibson assembly (New England Biolabs, E2611) and validated by DNA sequencing.

## Protein expression and purification

FreeStyle 293-F cells were maintained in FreeStyle 293 expression medium and transfected at a density of 1.8 x 10^6^ cells/ml. For 1 L of cells, 1 mg plasmid encoding the protein of interest was mixed with 3 ml of PEI MAX (Polysciences, 24765-1) in 40 ml fresh expression medium and incubated for 20 minutes at room temperature before transfection. Cells were harvested 72 hours after transfection. The cadherin-catenin complex was co-expressed by co-transfecting plasmids featuring GFP-tagged E-cadherin, untagged β-catenin, and untagged αE-catenin at a mass ratio of 1:2:2. The cadherin-catenin– vinculin complex was co-expressed by co-transfecting plasmids of Flag-tagged vinculin, GFP or GFP and Halo-tagged E-cadherin, untagged β-catenin, and untagged αE-catenin at a mass ratio of 1:2:4:4. Afadin constructs were expressed individually, and full pentameric complexes were reconstituted by mixing cadherin-catenin complexes with afadin fragments. Vinculin constructs were also expressed and purified alone for comparative binding assays.

Cells were lysed in Lysis Buffer (50 mM Tris-HCl pH 8.0, 150 mM NaCl, 2 mM MgCl_2_, 0.2% CHAPS, 2 mM β-mercaptoethanol, 5 mM ATP, 1 mM PMSF, 1 μg/ml aprotinin, leupeptin, and pepstatin). β-mercaptoethanol was excluded from Lysis Buffer for Flag affinity purification. Cell debris was removed by centrifugation at 20,000g for 30 minutes at 4 °C. GFP-tagged cadherin-catenin complexes, afadin constructs, and vinculin were affinity purified from supernatants using an anti-GFP nanobody coupled to NHS-activated Sepharose 4 Fast Flow resin (anti-GFP beads)^2^. The cadherin-catenin–vinculin complex was first affinity purified from supernatant using Flag M2 beads (Sigma, A2220). The bound complex was eluted with 200 μg/ml Flag peptide (Sigma, F3290), then further affinity purified using anti-GFP beads. The tags for all affinity-purified proteins were removed by 0.1 mg/ml TEV protease digestion^3^, then the cleaved proteins were eluted using Elution Buffer (25 mM Tris-HCl pH 8.0, 150 mM NaCl, 0.1% CHAPS). Afadin, vinculin, and the cadherin-catenin complex were further purified by anion exchange chromatography on a MonoQ 5/50 GL column (GE Healthcare), followed by size exclusion chromatography on a Superdex 200 increase 10/300 GL column (GE Healthcare). The cadherin-catenin–vinculin complex was directly polished by size exclusion chromatography on a Superose 6 10/300 column (GE Healthcare) without initial purification by anion exchange. Size exclusion columns were equilibrated with Size Exclusion Buffer containing 10 mM Tris-HCl pH 8.0, 100 mM NaCl, and 3 mM DTT.

Chicken skeletal muscle actin was prepared as previously described^4^ and stored at 4 °C in G-Ca buffer: 2 mM Tris-HCl pH 8.0, 0.5 mM DTT, 0.1 mM CaCl_2_, 0.2mM ATP, 0.01% NaN_3_ at 4 °C. F-actin was polymerized by mixing monomeric actin (diluted in G-buffer: 2 mM Tris-HCl pH 8.0, 0.5 mM DTT, 0.2 mM ATP, 0.1 mM MgCl_2_) with KMEI buffer (50 mM KCl, 1 mM MgCl_2_, 1 mM EGTA, 10 mM imidazole pH 7.0) and incubated at room temperature for 1 hour.

## F-actin co-sedimentation assays

4 μM of the cadherin-catenin complexes, either alone or mixed with 6 μM afadin fragment, were diluted in KMEI buffer and pre-cleared by ultracentrifugation at 80,000 rpm (278,000 x g) in a TLA-100 rotor (Beckman Coulter) for 15 minutes at 4℃. 50 μl of the supernatant was mixed with an equal volume of 10 μM F-actin. The mixture was incubated for 30 minutes at room temperature, then ultracentrifuged at 80,000 rpm (278,000 x g) for 30 minutes at 4℃. 40 μl of the supernatant was mixed with 10 μl of 5x SDS-PAGE Sample Buffer (50 mM Tris-HCl pH 8.0, 50 mM NaCl, 2% SDS, 0.025% Bromophenol Blue, 10% Glycerol, 5% β-mercaptoethanol). The pellet was washed twice with F-actin buffer (KMEI + 0.5 mM DTT) and dissolved in 125 μl of 1x SDS-PAGE Sample Buffer. Samples were separated by SDS-PAGE, stained with Coomasie Brilliant Blue, scanned on a LI-COR Odyssey scanner (LI-COR), and quantified with FIJI^5^.

## Pull-down assays

All proteins were expressed in FreeStyle 293-F cells. Flag-tagged afadin fragments were first bound to Flag M2 beads by affinity purification as described above. Cells expressing GFP-tagged individual domains of αE-catenin or the cadherin-catenin-vinculin complex were lysed in Lysis Buffer and spun at 20,000 x g for 30 minutes at 4 °C. The supernatants were collected and then incubated with afadin-bound Flag M2 beads. After 30 minutes of incubation, the beads were washed three times with Lysis Buffer, dissolved in 50 μl 1x SDS-PAGE Sample Buffer and analyzed by SDS-PAGE as described above.

## Stable cell line generation

Lentiviruses were produced in HEK293T cells by co-transfecting the pCDH lentiviral vector with the packaging plasmids PsPAX2 and pMG2.g at a mass ratio of 4:3:1 using lipofectamine 3000 (Thermo Fisher, L3000008). 48 hours after transfection, the culture medium containing the virus was harvested and filtered through a 0.45 μm pore size filter. 5 μg/ml polybrene (Sigma, TR-1003-G) was added to enhance infection efficiency. Afadin knockout EpH4 cells were infected with an equal amount of lentiviruses encoding Flag-tagged afadin constructs and rtTA. 48 hours after infection, cells were selected with 2 μg/ml of puromycin (Thermo Fisher, A1113803) and 800 μg/ml G-418 (Sigma, 108321-42-2) for 7 days. Surviving cells were diluted for single clone selection. Afadin expression was induced by addition of 0.05 μg/ml doxycycline (Sigma, D9891) to single-cell derived clones. Cells were analyzed by western blot or immunostaining 24 hours after induction.

## Immunofluorescence imaging

EpH4 cells at 90% confluency were fixed with 4% formaldehyde (Thermo Fisher, 28908) in Cytoskeleton Buffer (CB: 100 mM MES pH 6.1, 1.38 M KCl, 30 mM MgCl_2_, 20 mM EGTA) for 15 minutes, permeabilized in 0.1% Triton X-100 in CB for 10 minutes, then incubated with 100 mM glycine in CB for 10 minutes. After washing with DPBS (Gibco, 14190-144), cells were blocked in 2% Bovine Serum Albumin (BSA, Gemini Bio-Products, 700-101P) in DPBS for 1 hour at room temperature.

Blocked cells were incubated with primary antibodies in DPBS + 2% BSA overnight at 4℃. After washing with DPBS, cells were incubated with secondary antibodies at room temperature for 1 hour and washed with DPBS three times. The following antibodies were used for immunostaining: mouse anti-FLAG mAb (1:200; Sigma, F3165); rat anti-E-cadherin mAb (1:500, ThermoFisher, 13-1900); mouse anti-afadin mAb (1:200; Santa Cruz, sc-74433); Alexa Fluor 488 goat anti-mouse (1:250; Thermo Fisher, A11029); Alexa Fluor 488 goat anti-rabbit (1:250; Thermo Fisher, A32731); Alexa Fluor 647 donkey anti-rat (1:250; Jackson ImmunoResearch, 712-605-153). Alexa Fluor 568 Phalloidin (1:40; Thermo Fisher, A12380) was used to visualize F-actin. Image series with a depth range of 1.2 μm and step size of 0.2 μm along the z axis were acquired on a Nikon Ti-E microscope equipped with a CFI Apo 100x oil immersion objective (numerical aperture 1.49). Epifluorescence illumination was provided by a LED illuminator (Lumencor). Images were acquired on a Zyla 4.2 sCMOS camera (Andor) and quantified using FIJI^6^.

## Western blotting

Western blots were performed following standard protocols. EpH4 cells were washed with DPBS once and lysed in 1x SDS-PAGE Sample Buffer. Samples were separated by SDS-PAGE and transferred onto a nitrocellulose membrane (Bio-Rad, 1620115). The membranes were blocked with SuperBlock Blocking Buffer (Thermo Fisher, 37537) and subsequently incubated with primary antibodies and secondary antibodies, both diluted in SuperBlock T20 Blocking Buffer (Thermo Fisher, 37536). The membranes were scanned on a LI-COR instrument. The following antibodies were employed: mouse anti-FLAG mAb (1:1000; Sigma, F3165); rabbit anti-afadin mAb (1:1000; Thermo Fisher, 700193); mouse anti-GAPDH mAb (1:2000; abcam, ab8245); IRDye 800CW goat anti-mouse (1:15000; LI-COR, 926-32210); IRDye 800CW goat anti-rabbit (1:15000; LI-COR, 926-32211).

## TIRF microscopy clustering assay and image analysis

No. 1.5 24 x 60 mm glass coverslips were cleaned by sequential sonication in 100% acetone for 30 minutes, 100% ethanol for 10 minutes, and 2% Hellmanex for 120 minutes. After rinsing with MilliQ water, the coverslips were functionalized by incubating with 0.9 mg/ml mPEG-silane (Laysan Bio, MPEG-SIL-5000), 0.1 mg/ml biotinated mPEG-silane (Laysan Bio, Biotin-PEG-SIL-5K), 10 mM HCl, and 96% ethanol overnight. Subsequently, the coverslips were rinsed with 100% ethanol and MilliQ water, air dried, and stored in a container filled with nitrogen gas.

F-actin was prepared by co-polymerizing 0.72 μM Atto488-labelled and 0.08 μM biotin-labelled monomeric actin (Hypermol, 8153-02 and 8109-01, both diluted in G-buffer) at room temperature for 60 minutes. The Halo-tagged tetrameric Eβα(CA)-Vinculin(T12-IV) complex was fluorescently labelled with Janelia Fluor 549 HaloTag ligand (Promega, GA1110) at a molar ratio of 1:2 at room temperature for 4 hours. After removing free dye using a Pierce dye removal column (Thermo-Fisher, 22858), the labelled complex was diluted to 4 μM in Imaging Buffer, consisting of Motility Buffer (MB: 20 mM MOPS pH 7.4, 5 mM MgCl_2_, 0.1 mM EGTA, 50 mM NaCl, 1 mM DTT), supplemented with 15 mM glucose, 100 μg/mL glucose oxidase, and 20 μg/ml catalase. Prior to imaging, the labelled complex was precleared by ultracentrifugation at 80,000 rpm (278,000 x g) for 12 minutes at 4℃ in a TLA100 rotor.

Imaging wells were assembled by affixing a CultureWell gasket (GraceBio, 103280, 1 mm in thickness and 6 mm in diameter) onto the cover slip surface. After blocking with 1 mg/mL casein (Sigma, C0406) in MB for 1 hour, the well was incubated with 1 μM streptavidin (Sigma, 189730) in MB for 2 minutes, rinsed with MB, coated with labelled F-actin for 1 minute, and rinsed again with MB. The labelled tetramer, either alone or with dark afadin-CC at 4 times the tetramer’s concentration, was introduced to replace MB. The well was either immediately mounted for time-lapse imaging or placed in a humidified container for 10 minutes before individual snapshot imaging.

Dual-color Total Internal Reflection Fluorescence (TIRF) imaging was conducted using a Nikon Ti-E microscope equipped with an H-TIRF motorized module and NIS-Elements software (Nikon). F-actin and the Halo-tagged tetrameric complex were excited by 488 nm and 561 nm lasers (Agilent), respectively. For time-lapse imaging, frames were acquired every 2 seconds using a CFI Apo 100X TIRF oil immersion objective (numerical aperture 1.49), a quad filter (Chroma), and an iXon EMCCD camera (Andor). Individual snap shots were acquired with identical exposure settings.

Image analysis was performed with custom Python scripts using functions from the scikit-image package^7^. Uneven illumination in the tetramer channel was corrected using rolling-ball background subtraction with a ball radius of 100 pixels. F-actin and tetramer masks were generated by thresholding (Yen method) and binarizing the corresponding channels. A mask corresponding to tetramer specifically bound to F-actin (“bound tetramer mask”) was then obtained through the logical conjunction of the actin and tetramer masks. The tetramer bound fraction of F-actin was calculated by dividing the area of the bound tetramer mask by the area of the total F-actin mask. The fraction bound was then plotted versus tetramer concentration and fit by the Hill equation^8^.

## Cryo-EM sample preparation

2.5 μM of the tetrameric Eβα(CA)-Vinculin(T12-IV) and 5 μM of afadin-CC were premixed before use. 3 μl of 0.6 μM F-actin (polymerized in the presence of 0.01% Nonidet P40 (NP40) substitute (Roche) to reduce ice thickness) was applied onto a freshly plasma cleaned C-flat 1.2/1.3 holey carbon Au 300 mesh grid (Electron Microscopy Sciences) in a Leica EM GP plunge freezing apparatus operating at 25°C and 100% humidity. After 1 minute incubation, 3 μl of the premixed protein complex was added and mixed thoroughly with the F-actin drop. After an additional 2 minute incubation, the grid was blotted from the back with Whatman no. 5 filter paper for 5 seconds and plunge-frozen in liquid ethane.

## Cryo-EM and cryo-ET data acquisition

Cryo-EM data were acquired on a Thermo Fisher Titan Krios transmission electron microscope operating at 300 kV equipped with a Gatan K2-summit detector using SerialEM^9^. All image stacks were recorded at a nominal magnification of 29,000X in super-resolution mode with a calibrated pixel size of 1.03 Å at the specimen level (super-resolution pixel size of 0.515 Å / pixel). Each exposure was dose fractionated across 40 frames with a total electron dose of 61 e^-^ / Å^2^ (1.53 e^-^ / Å^2^ / frame) and a total exposure time of 10 s. The target defocus values ranged from –0.8 to –2.0 μm. Exposures were collected using the beam tilt / image shift strategy, targeting 9 holes per stage translation.

Cryo-ET data were obtained on a spherical-aberration (Cs) corrected Titan Krios transmission electron microscope operating at 300 kV equipped with a Gatan K3 direct electron detector and a BioQuantum energy filter (slit width 20 eV). The tilt series was recorded using a dose-symmetric scheme^10^ from –60° to 60° with a tilt increment of 3°. Exposures at each tilt angle were acquired at a nominal magnification of 26,000X, corresponding to a calibrated pixel size of 2.6 Å at the specimen level (super-resolution pixel size of 1.3 Å / pixel). Each exposure was fractionated into 12 frames (0.22 e^-^ / Å^2^ / frame) across an exposure time of 0.6 s, with a total electron dose across the tilt series of 107.82 e^-^ / Å^2^.

The tilt series was collected at a target defocus value of –3.5 μm.

## Cryo-EM image processing

11,061 movie stacks were motion corrected with a binning factor of 2 (resulting in a 1.03 Å pixel size) using MotionCor2^11^. Contrast Transfer Function (CTF) estimation was carried out with CTFFIND4^12^. The signal from bound cadherin-catenin-vinculin complexes prevented standard filament pickers from accurately tracing the filament center. Therefore, particle picking was conducted using a customized neural-network–based approach, adapting our previously established approach for processing actin filaments^13^. Specifically, a library of 312 synthetic volumes was created featuring actin filaments that were either bent or straight. These filaments had varying decoration patterns of a coarse cadherin-catenin complex model which we constructed by aligning crystal structures of individual components (PDB IDs: 1I7X, 1DOW, 4IGG)^14–16^. To encompass plausible cadherin-catenin complex decoration patterns, three potential orientations of the complex relative to F-actin were generated, and for each complex orientation and F-actin curvature, a set of filaments was generated with one bare filament, one fully decorated filament, and partially decorated filaments with random occupancies. This library was used to train a neural network that successfully segmented F-actin at the exclusion of neighboring signal from the flexibly bound cadherin-catenin complex (Figure S3). Filament splines were traced through these segmented micrographs to generate initial picks.

1,524,620 picked coordinates were imported into RELION-3.1^17^ and extracted with a box size of 448 pixels, then binned by 2. The extracted particles were imported into Cryosparc v4.2^18^ for reference-free 2D classification. 521,858 particles from high-quality 2D classes were selected for *ab initio* reconstruction and masked homogeneous refinement. The refined particles were imported back into RELION-3.1, then symmetry expanded with a helical twist of –167°, rise of 27 Å, and 23 asymmetric actin subunits. This effectively captured nearly all the filaments in the dataset. After duplicate removal, 6,729,146 particles were re-extracted with a box size of 448 pixels, binned by 4, and subjected to reference-free 2D classification. High quality 2D classes displaying additional densities along both strands of the filament were selected and subjected to unsupervised 3D classification while applying helical symmetry. Particles with clear filament decoration were refined and symmetry-expanded by applying a helical twist of –167° and a rise of 27 Å across 3 asymmetric actin subunits. After duplicate removal, particles underwent multiple rounds of masked 3D classification without image alignment.

Particles exhibiting clear afadin coiled-coil density (470,416 particles) were re-centered on residue K889 of the αE-catenin ABD, re-extracted with a box size of 448 pixels without binning, and subjected to CTF refinement, Bayesian polishing, and masked 3D auto-refinement, resulting in a postprocessed density map at 2.9 Å resolution. To enhance the resolution of the αE-catenin CTE, particles underwent another round of focused 3D classification with a tight mask covering only the densities of the CTE and afadin’s coiled-coil. One high-quality class with 99,745 particles was selected and refined to a resolution of 3.1 Å.

2D classes displaying additional densities along one strand of the filament were re-extracted with a box size of 768 pixels and then binned by 4. Following unsupervised 3D classification, particles showing the best occupancy of additional densities were re-extracted with a box size of 448 pixels and subjected to masked 3D auto-refinement by applying helical symmetry. The refined particles were symmetry expanded by applying a helical twist of 26° and rise of 55 Å across 3 actin subunits. After duplicate removal, particles were re-extracted with a box size of 512 pixels and subjected to masked 3D auto-refinement without applying helical symmetry, resulting in a density map at 4.4 Å resolution. After one round of supervised 3D classification, the single class with high afadin-CC occupancy was selected (44,548 particles) for subsequent CTF refinement, Bayesian polishing and masked 3D auto-refinement, yielding a postprocessed density map at 4.1 Å resolution. The refined particles were used for continuous conformational variability analysis using CryoDRGN^19^.

## Cryo-ET reconstruction

Individual frame motion correction and CTF estimation of summed tilt exposures were performed in WARP^20^. The tomogram was reconstructed using the IMOD software package^21^. The tilt series was binned by 3 to a final pixel size of 7.8 Å and aligned using the patch-tracking approach. The tomogram was reconstructed using a SIRT-like filter, equivalent to 8 iterations.

## Model building and refinement

The models of αE-catenin ABD bound F-actin (PDB: 6UPV) and the coiled-coil of afadin predicted by AlphaFold^22^ were rigid body docked to the 3.1 Å resolution density map in UCSF Chimera^23^. Subsequently, the complete model was manually built and adjusted in Coot, flexibly fitted with ISOLDE in UCSF ChimeraX^24,25^, refined using Phenix.Real_space_refine, and validated with MolProbity as implemented in Phenix^26,27^.

## Local helical parameter measurements

Filament twist and rise were measured using a custom Python script described previously^28^. In brief, a model containing 15 actin subunits was built by rigid-body fitting each individual actin subunit (PDB:7R8V, chain B) into the density map of each analyzed filament reconstruction in Chimera. Three copies of each model were iteratively aligned on the four terminal actin subunits at the pointed end. After deleting the overlapping subunits from the two end copies, an extended filament model with 37 subunits was created and used for twist and rise measurement. To avoid edge effects, only the twist and rise values of the central 10 subunits were retained for further analysis.

## Occupancy measurements

Atomic models of one actin subunit with bound ⍺E-catenin–ABD and afadin-CC were individually fit into each reconstructed frame of the first principal component of the cryoDRGN variability trajectory. Fitting for sites with low or no ⍺E-catenin–ABD and afadin-CC density was driven by the actin density. Density for the filament and the bound complex were separated using the “split map” command in Chimera. The integrated intensity of each segmented bound complex sub-volume was computed, then normalized to the site with the highest integrated intensity. As the bound ⍺E-catenin– ABD and afadin-CC at this site had approximately the same intensity as F-actin, this measurement serves as a proxy for occupancy. The mean normalized intensity for each strand of each reconstructed frame from the cryoDRGN trajectory was then computed.

## Plotting and analysis

Figures and movies were generated with UCSF ChimeraX^24^ and FIJI. All statistical analysis and plotting were performed in GraphPad Prism. Multiple sequence alignments were performed with Clustal Omega^29^ and rendered with Jalview^30^.

